# EM and component-wise boosting for Hidden Markov Models: a machine-learning approach to capture-recapture

**DOI:** 10.1101/052266

**Authors:** Robert W. Rankin

## Abstract

This study introduces statistical boosting for capture-mark-recapture (CMR) models. It is a shrinkage estimator that constrains the complexity of a CMR model in order to promote automatic variable-selection and avoid over-fitting. I discuss the philosophical similarities between boosting and AIC model-selection, and show through simulations that a boosted Cormack-Jolly-Seber model often out-performs AICc methods, in terms of estimating survival and abundance, yet yields qualitatively similar estimates. This new boosted CMR framework is highly extensible and could provide a rich, unified framework for addressing many topics in CMR, such as non-linear effects (splines and CART-like trees), individual-heterogeneity, and spatial components.

## 1. Introduction

Multi-model inference (MMI) has become an integral part of the capture-mark-recapture (CMR) literature. By CMR, I refer to the survey design and statistical modelling of abundance and survival of marked animals under imperfect detection, using individual time-series of recaptures. By MMI, I loosely refer to a variety of strategies such as model-selection, model-averaging, and regularization techniques such as shrinkage estimators (e.g. some random-effects models; Royle & Link, 2002) and sparse estimators. A good overview is by Leeb & Pötscher (2009). These strategies may be used to address research goals such as: finding ecologically important covariates; deciding which model-cum-hypothesis has most support; incorporating “model uncertainty” into estimates; or seeking parsimony in estimation, such as estimating survival across sex and age classes, and doing so without over-fitting.

Among these related goals, we may categorize them into two distinct objectives: estimation/prediction vs. selection of the “correct” model or “best approximating” model. Often, these two objectives cannot be achieved by the same MMI procedure (Shao, 1993; Yang, 2005; Leeb & Pötscher, 2005; Vrieze, 2012; Aho et al., 2014). Estimation is generally heralded by shrinkage-estimators and the Akaike Information Criterion (AIC; Akaike, 1998, 1974), whereas selection is championed by sparse-estimators and the Schwarz-Bayes Criterion (BIC; Schwarz, 1978). This paper will introduce a new MMI technique for capture-mark-recapture called “boosting”, and I will show how it fits into the two domains of MMI.

Boosting is a type of shrinkage estimator, a class of techniques that (crudely) achieve the goals of MMI with a single smoothing model. Crucially, model complexity can “shrink” along a continuum, in contrast to all-subsets model-selection where there is a discrete set of fixed-effect models with different numbers of parameters. Shrinkage estimators were first motivated by Royle & Link (2002) for CMR, in which case they advocated for a random-effects Bayesian model. In this paper, I present a new boosting algorithm, which could be considered as the Frequentist answer to Royle & Link.

To understand shrinkage, consider the classic example of survival and its fixed-effects extremes: time-varying survival vs. time-constant survival. In CMR notation, these are known as *ϕ*(*t*) and *ϕ*(·), respectively. The former is difficult to reliably estimate, whereas the latter is often a poor reflection of reality. Shrinkage estimators will achieve an intermediate solution between the two extremes. In other words, *ϕ*(*t*) is shrunk towards *ϕ*(·).

The question then becomes, how much shrinkage? To Bayesians, like Royle & Link (2002), the answer is to use prior distributions. To a Frequentist, the amount of shrinkage is decided by prediction error: we find a model that can both explain the observed data and make good predictions on new data. CMR practitioners may not think of themselves as seeking models with good predictive performance, but their tool of choice, the AIC/c, is based on a predictive error called the KL-loss (Akaike, 1974, 1998). Likewise, boosting methods are highly *efficient* at minimizing prediction error and estimation error (Bühlmann & Yu, 2003; Meir & Rätsch, 2003). This makes boosting very philosophically similar to model-selection by AIC (Leeb & Pötscher, 2009). Therefore, boosting should be of great interest to CMR practitioners who are already using the AIC for model building.

However, boosting can do things that AIC/c model-selection cannot. For example, it can include splines for non-linear effects (e.g., a non-linear change in survival with age). It can include classification and regression trees (CART; Hothorn et al., 2006) for automatic discovery of higher order interactions (such as a three way interaction of sex, time, and age on capture-probability). It can include spatial effects (Kneib et al., 2009; Tyne et al., 2015). It can deal with “high-dimensional” covariate data, such as sorting through dozens or hundreds of potential environmental variables, even under small sample sizes. It also does a better job of handling “model uncertainty” under the scourge of multi-collinearity (Mayr et al., 2014), which troubles the model-averaging approach (Cade, 2015). Boosting is also related to many other types of popular techniques, such as being a type of Generalized Additive Model (Schmid et al., 2010; Hofner et al., 2014) and *𝓁*_1_-regularization (a.k.a. the Lasso; Bühlmann & Yu, 2003; Efron et al., 2004; Tibshirani, 2011). This versatility has led some to call boosting the “unified framework for constrained regression” (Hofner et al., 2014). This paper introduces this powerful framework to CMR.

Many of the above benefits should interest CMR practitioners (especially believers of the AIC approach). Perhaps most importantly, boosting excels in one particular domain which is terribly onerous for all-subsets model-selection: the scourge of high-dimensionality. Every additional covariate leads to an exponential increase in the number of possible fixed-effect models. This is due to the multi-parameter nature of CMR models: we must perform model-selection on both the survival parameter as well as the capture parameter. In this paper, I will consider an example with just three covariates (sex, time, and an environmental covariate) which results in 64 fixed-effects models. With a fourth and fifth covariate, the number of fixed-effect models would explode to 196 and 900, respectively. This computational burden is quickly prohibitive for all-subsets model-selection, with even a small number of covariates. Consequentially, some recent CMR studies using AIC/c model-selection have taken computational shortcuts, such as step-wise selection (Pérez-Jorge et al., 2016; Taylor et al., 2016), an out-dated procedure that is strongly discouraged for many reasons (Burnham et al., 2011). In contrast, boosting can sort through all covariates and their interactions in just one model, because covariate selection is integrated within the fitting procedure.

I will introduce CMR boosting for the two-parameter open-population Cormack-Jolly-Seber model (CJS; Cormack, 1964; Jolly, 1965; Seber, 1965), for estimating survival and abundance under imperfect detection. The simplicity of the CJS will suffice to prove the new boosting algorithm for CMR data; such data is not possible to analyze using conventional boosting algorithms. Conventional boosting methods assume independent data-points in order to perform gradient descent (i.e., step-wise minimization of a loss function), whereas CMR capture-histories consist of serially-dependent observations. The key innovation of this paper is to garner conditionally independent observations by imputing time-series of latent states, a routine trick from Hidden Markov Models (HMM). In CJSboost, we alternate between boosting the parameters (conditional on latent states) and imputing expectations of the latent states (conditional on the parameters), and repeating *ad infinitum*. I will prove this framework on the simple and manageable CJS model, with the ultimate goal to refine the method on more complex models, such as POPAN and the Robust Design and spatial capture-recapture.

By focusing on a simple CJS model, I will also elucidate some of the technical challenges and limitations of boosting. The most obvious challenge is the computational burden of multiple cross-validation steps. Another less obvious limitation is that boosting is generally unsuitable for making inferences about the “true model” or discriminating among truly influential covariates vs non-influential covariates, i.e., it is not model-selection *consistent*. This is true for all procedures that are optimized for prediction/estimation, including the AIC/c (Yang, 2005; Leeb & Pötscher, 2009; Vrieze, 2012; Aho et al., 2014). These loss-efficient procedures have a well-known tendency to prefer slightly more-complex models (Shao, 1997) and they can result in false discoveries when misused to find the “true model”. As a possible remedy, I suggest combining CJSboosting with a new regularization-resampling technique called stability selection (Meinshausen & Bühlmann, 2010) to make inferences about which covariates are truly influential. Therefore, CMR practitioners can use CJSboost for either efficient estimation or consistent model-selection/model-identification.

First, I will provide some background theory about model-selection and shrinkage, as well as a brief introduction to conventional boosting algorithms. Then, I will use simulations and a classic dataset (Lebreton et al., 1992) to illustrate CJSboost and benchmark it to AICc model-selection and model-averaging. Finally, I will end with a simulation that is computationally impossible for AICc-based inference: model-selection of a CJS model with 21 covariates. This is unheard of in CMR, until now.

For R code (R Core Team, 2016) and a tutorial, see the online content at http://github.com/faraway1nspace/HMMboost/.

## 2. Methods

### 2.1. Background

#### 2.1.1. Capture-Recapture and the Cormack-Jolly-Seber Model

Imagine that we wish to study the abundance and survival of an open-population of animals. At regular time-intervals *t ∈* {*1, 2, 3*, …, *T*}, we randomly capture, mark, and release individual animals. In subsequent *t ≥* 2, we recapture some of these already-marked animals with probability *p*_*i,t*_, conditional on an animal being alive at *t*. Animals may die between capture periods *t*–1 and *t*, or survive with probability *ϕ*_*i,t*_. Recaptures are scored as the binary outcome *y*_*i,t*_ *∈* {0, 1} for {*no-capture,re-capture*}. **y**_*i*_ is the time-series of captures for individual *i*, called a *capture-history*. The ragged matrix **Y**^(*n×T*)^ includes the capture-histories of all *n* unique individuals who were observed.

Our goals are two-fold: i) to estimate the abundance of marked animals *N*_*t*_ for each capture period *t >* 1; and ii) estimate survival *ϕ*, including its sources of variation, such as temporal variation or individual variation. The above formulation is the Cormack-Jolly-Seber open population model (Cormack, 1964; Jolly, 1965; Seber, 1965). We can estimate the parameters *p*_*i,t*_ and *ϕ*_*i,t*_ by maximizing the CJS likelihood:

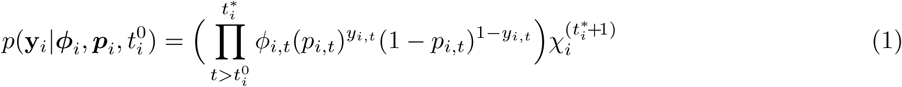

where 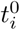 is the capture-period in which individual *i* was first captured; 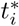 is the capture-period when individual *i* was last observed; and 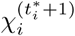 is the probability of never being seen again after 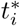 until the end of the study, 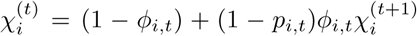. Notice that 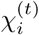 is calculated recursively. Given 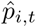, we can estimate the abundance of animals at time *t* using a Horvitz-Thompson-type estimator: 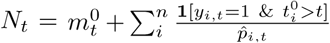, where 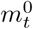 is the number of animals whose first capture was at time *t* (McDonald & Amstrup, 2001).

A key point is that the captures are serially-dependent and cannot be considered independent; in other words, the CJS likelihood (1) is evaluated on an entire capture-history, *not* per capture. This is mathematically embodied by the recursive term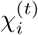. For gradient descent algorithms, like boosting, we require independent data points. If we reformulate the CMR system as a HMM, we can garner conditional independence through the use of latent states *z*_*i,t*_ *∈* {0, 1} to represent {dead, alive}. When *z*_*i,t*_ = 1, then individual *i* is alive and available for capture at time *t*, and the probability of a capture is simply *p*(*y*_*i,t*_ = 1*|z*_*i,t*_ = 1) = *p*_*i,t*_. However, if *z*_*i,t*_ = 0 then individual *i* is dead and unavailable for capture at time *t*; therefore the probability of a capture is zero.

The use of latent states and boosting is not new (Ward et al., 2009; Hutchinson et al., 2011). The novelty of the CJSboost approach is that the latent states obey Markovian transition rules and form a serially-dependent time-series. For example, a trailing sequence of no-captures **y**_*t*:*T*_ = [0*, …,* 0]^T^ has many possible state-sequences, but once *z*_*t*_ = *dead* then also *z*_*t*+1_ must equal *dead*. Fortunately, we can utilize well-developed HMM tools to estimate all the permissible state-sequences **z**. This is a key point which will be developed further when I describe the CJSboost algorithm.

#### 2.1.2. Prediction, Estimation and Generalization Error

There are many types of MMI techniques that share an implicit property of making optimal predictions. This is true for shrinkage estimators, like boosting, and the AIC and their cousins (i.e. what Aho et al., 2014, called “A-type” thinking). Here, prediction has a more technical meaning than, e.g., the layman idea of weather forecasting or predicting the next USA president. It means that if we collect a new sample of data *y*^(new)^ from the population 𝕐, our predictive model should be able to accurately estimate the *y*^(new)^ values. More formally, we wish to minimize the error in predicting *y*^(new)^, for all theoretical data-sets that we might randomly sample from the population distribution of 𝕐. Notice that this predictive framework is not explicitly about testing hypotheses nor accurate estimation of parameters, but it nonetheless serves as a principled means of model-building: we desire a model that is complex enough to fit to the observed data and make good predictions on new data, but does not over-fit the observed data. This is one way to codify *parsimony*.

We can formalize this intuition as the following. Consider that we have a family of models G which map covariate information 𝕏 to the response variable, i.e., *G*: 𝕏 *→* 𝕐. Our sample of data 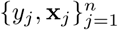 arises from the unknown population distribution *P*. The optimal model *G* is that which minimizes the following *generalization error*:

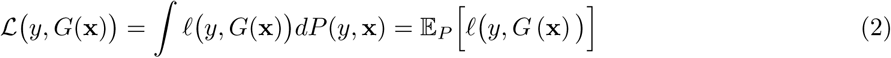

where *𝓁* is a *loss* function: it scores how badly we are estimating *y* from *G*(**x**). *ℒ* is the *expected loss*, a.k.a, the *risk* (Bühlmann & Yu, 2003; Meir & Rätsch, 2003; Murphy, 2012a). Here, the integral is just a mathematical way of saying that we are minimizing the loss over the entire theoretical population, and any new samples from this population.

There are many types of loss-functions. Akaike (1998) makes the case for using the (negative) log-Likelihood; in which case Eqn. 2 becomes equivalent to minimizing the Expected (negative) log-Likelihood (which is not to be confused with Maximum Likelihood Estimation). In fact, the Expected log-Likelihood is seen in Eqn 1.1 of Akaike’s seminal derivation of the AIC (Akaike, 1998). This emphasizes the fundamental similarity between the AIC and any estimator that minimizes (2).

While minimizing the expected loss is ostensibly about predicting new values of the response variable *y*, it also has desirable properties for estimation. This is crucially important because CMR practitioners are not interested in making predictions about new capture-histories. Instead, we want to minimize the error of estimating abundance and survival. Fortunately, as noted by Akaike (1974), minimizing the Expected (negative) log-Likelihood is *efficient*. This means that by minimizing the expected loss (2) we also minimize the square-error between the estimated model parameters and their true values. This connection is straight-forward in multiple linear regression (Copas, 1997), but may only be approximately true for capture-mark-recapture. Through simulations, I will explore this estimation error for the CJS and the AICc (sections 2.4).

#### 2.1.3 Regularization and shrinkage

One cannot measure the expected loss or generalization error (2); it requires having data for the entire population. Instead, we are forced to work only with our sample of data, and proceed to minimize the *empirical risk*:

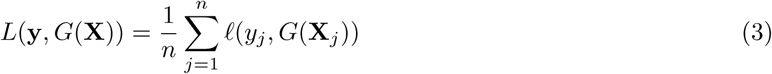

The difference between (2) and (3) is that the former integrates the loss over the entire population, while the latter only calculates the loss on the observed data. Minimizing the empirical risk is easy. In when the loss function is the negative log-likelihood, it is the Maximum Likelihood solution. But, at finite sample sizes, it tends to *over-fit* a sample, make bad predictions, and have higher estimation errors (Copas, 1983, 1997).

The question then becomes: how can we minimize something we cannot see (the generalization error), when all we have to work with is the observed data and empirical risk? Akaike (1998, 1974) answered this question with the AIC, which was to approximate the Expected (negative) log-Likelihood with 2*L*(**y***, G*(**X**))+ 2*‖G‖*_*o*_, where the second term is the number of parameters in *G*, a.k.a the *𝓁*_0_ norm^1^. The approximation works well at large sample sizes for linear regression and auto-regressive models, but is less exact for CMR models.

Another answer comes from Learning Theory, called regularization. The theory tells us that if we constrain the complexity of our function space, we can use the same procedure that minimizes the empirical risk, but still bound the generalization error (Bühlmann & Yu, 2003; Meir & Rätsch, 2003; Mukherjee et al., 2003). Practically, this implies that we penalize the complexity of *G* and prevent the procedure from fully minimizing *L*. Popular examples are the Lasso (Efron et al., 2004; Tibshirani, 2011) and Ridge regression, which have penalties on the *𝓁*_1_-and *𝓁*_2_-norms, respectively; hence, they are known as *𝓁*_1_-and *𝓁*_2_-regularizers. Boosting is generally equivalent to *𝓁*_1_-regularization (under certain circumstances; Efron et al., 2004; Bühlmann & Hothorn, 2007).

In boosting, the principal means of regularization is by *functional gradient descent* and *early-stopping*. Gradient descent means: i) we start with a very simple model *G*^(0)^ that has a high empirical risk *L*^(0)^ ; and then ii) we take tiny steps that reduce *L* towards its global minimum, where each *m*^*th*^ step slightly increments the complexity of the model *G*^(*m*)^. If we run the gradient descent until *m → ∞*, we would minimize the empirical risk and get a *fully-saturated model G*^(*m→∞*)^, which is generally equivalent to Maximum Likelihood Estimation. But, we stop short at some *m*_stop_ *« ∞*. Figure 1 (*bottom panel*) shows the gradient of the empirical risk.

**Figure 1:**
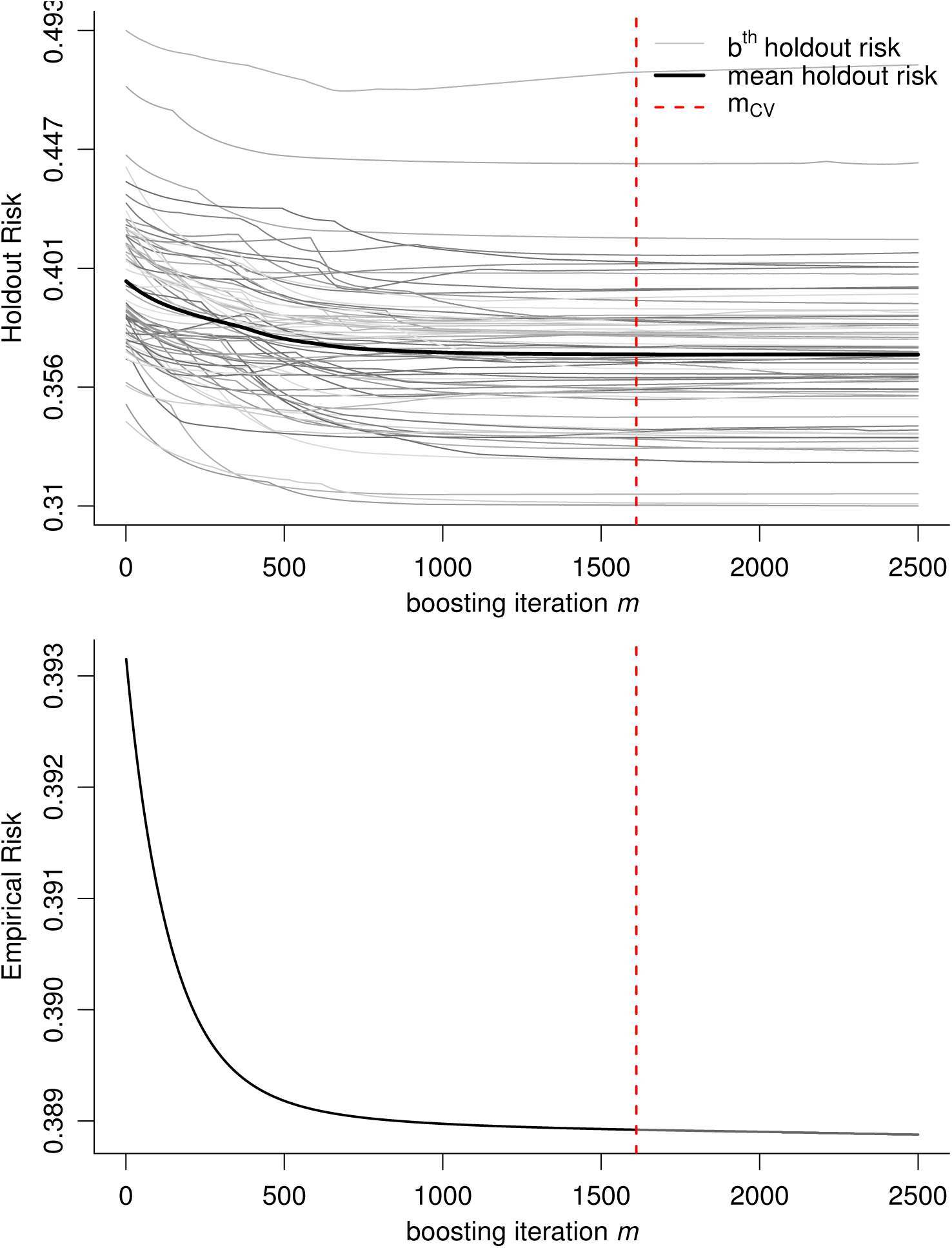
*Top*: Visualization of the step-wise minimization of *generalization error* by CJSboost (a.k.a the *expected loss* or *risk*), which is approximated by the mean holdout-risk (solid **black line**) from bootstrap-validation. Each *m* step along the x-axis is a boosting iteration, which adds one base-learner and increases the complexity of the model. At *m* = *m*_*cv*_ (red dashed line), the mean holdout-risk is minimized; beyond *m*cv the model is over-fitting. Each *b*^*th*^ gray line represents the holdout-risk predicted from one CJSboost model trained on a bootstrapped sample of capture-histories and then evaluating the holdout-risk on the out-of-sample data. *Bottom*: The empirical risk of the final statistical model using the full dataset. The model increases in complexity until it stops early at *m* = *m*_*cv*_. The empirical risk is the negative log-Likelihood of the Cormack-Jolly-Seber model. Running the algorithm for *m → ∞* will result in the MLE solution. The difference between the MLE model and the model at *m* = *m*_*cv*_ is *shrinkage*.

Why would we want to stop-short and not maximize the model-fit to the data? It turns out that, at finite sample sizes, the best predictors which minimize the generalization error have *shrinkage*: the estimates are shrunk away from the MLEs of the fully-saturated model and are pushed towards the simple model *G*^0^ (Copas, 1983, 1997). Optimal predictors are never as extreme as the MLEs. This predictive principle generally holds true for estimation as well; it was discovered as early as the 1950’s by Stein (1956) and James & Stein (1961). It was incendiary at the time because shrinkage estimators are *biased*. For example, Figure 2 compares true and estimated values from CJSboost, and I suspect most ecologists will find it alarming: it clearly shows the bias of shrinkage. A simple way to understand the optimality of shrinkage is through the idea of the “bias-variance trade-off”: we may be slightly biased but our estimates are likely to be closer to the truth (low-variance), whereas the MLEs are unbiased but may vary wildly with a new sample of data (high-variance). The Appendix E provides a primer about the bias-variance trade-off, and compares how CJSboost and AIC methods each negotiate this trade-off to minimize an expected loss.

**Figure 2:**
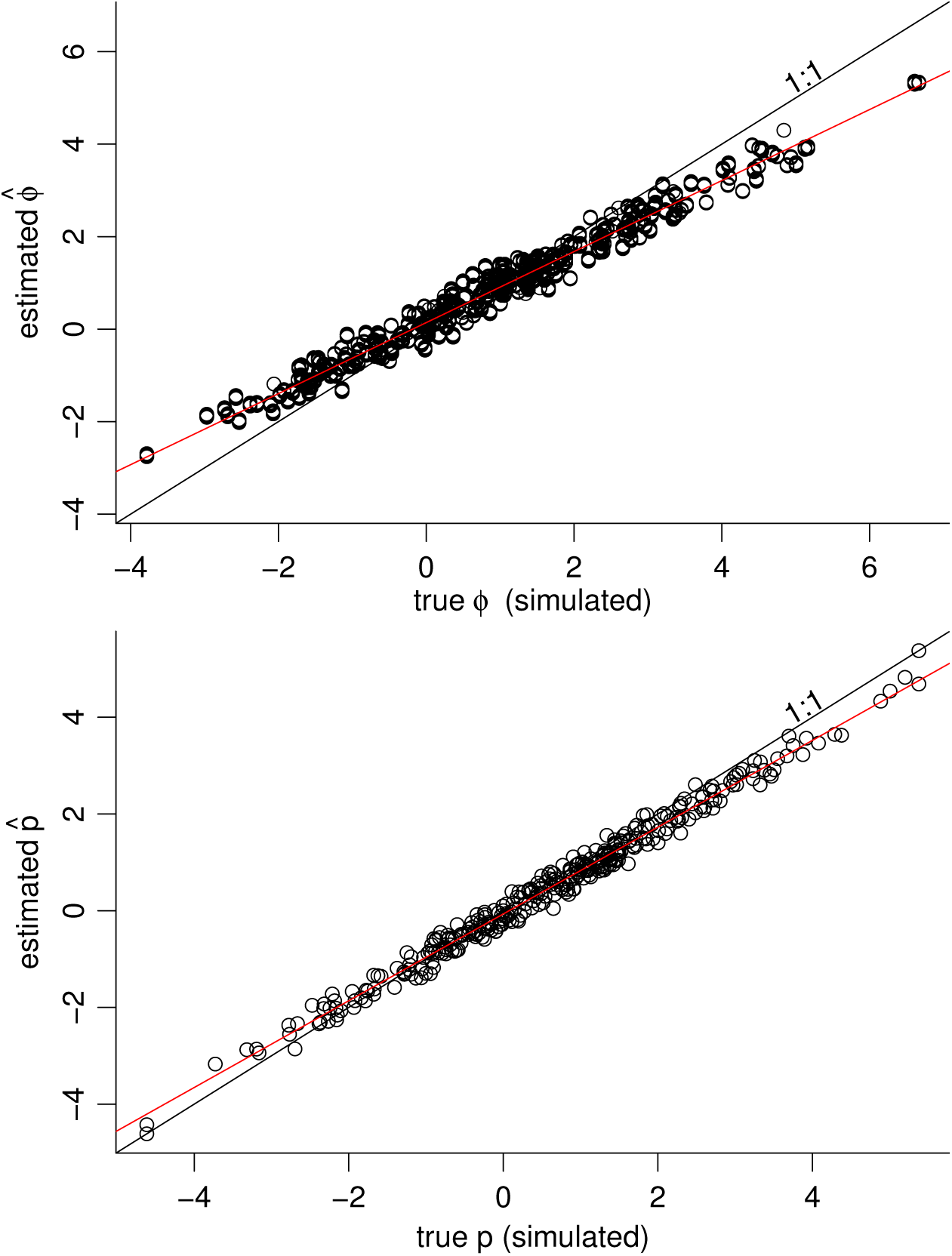
Visualization of shrinkage, by comparing the true (simulated) values of survival (*ϕ*_*i,t*_; *top*) and capture-probability (*p*_*i,t*_; *bottom*) vs. the CJSboost-EM estimates. Each point is an individual *i* at capture-period *t*. The CJSboost estimates have some downward bias (evident in the difference between the 1:1 line and the estimates’ **red** trend-line) due to shrinkage of coefficients to the intercept-only model. The amount of bias is our principle means of negotiating the “bias-variance trade-off” for optimal prediction.

Of course, we cannot measure the expected loss, so we must approximate it with the *average holdout-risk* using cross-validation or bootstrap-validation. We measure the empirical risk on out-of-sample subsets of bootstrapped data. The goal is to tweak the complexity of the model, by varying the regularization parameters, such that the average holdout-risk is minimized. Figure 1 (*top panel*) shows an example of minimizing the average holdout-risk at *m* = *m*_*CV*_. For a large number of bootstrap resamples, minimizing the average holdout risk will also minimize the expected loss. The wondrous utility of AIC is that it is generally equivalent to minimization of a leave-one-out cross-validation criteria (Stone, 1977; Shao, 1993, 1997).

#### 2.1.4. Introduction to boosting

The previous sections pertained generally to shrinkage estimators and MMI. I will now tie these ideas together with boosting before describing the CJSboost algorithm in section 2.2.1. This overview will focus only on the statistical view of boosting, whereas its full history and origins in machine-learning can be found in Meir & Rätsch (2003) and Mayr et al. (2014).

Statistical boosting can be thought of in two ways. One, it is an iterative method for obtaining a statistical model, *G*(*X*), via functional gradient descent (Breiman, 1998; Friedman et al., 2000; Friedman, 2001; Breiman, 1999; Schmid et al., 2010; Nikolay Robinzonov, 2013), where 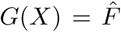 and 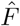 is the *fit-vector*, the expected values of *Y* based on covariate data *X*. Although boosting has origins in classification algorithms, we now know that it is equivalent to regularized regression, such as the Lasso (Bühlmann & Yu, 2003; Efron et al., 2004, under certain conditions).

Second, boosting is the step-wise construction of an ensemble model *𝒢*:= {*g*^(1)^*, g*^(2)^*, …, g*^(*m*)^}, composed of many weak prediction functions *g*, somewhat similar to model-averaging (Hand & Vinciotti, 2003). The prediction functions arise from base-learners *b*, which are any function that can take data (*x, y*) and make a predictor *g*(*x*) to predict *y* from 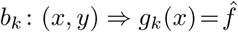. The fitting function *b* may be a Least-Squares estimator *b*_OLS_, or Penalized Least-Squares estimator *b*_PLS_, or recursive-partitioning trees *b*_trees_ (a.k.a CART), or low-rank splines *b*_spline_, or many others. The variety of base-learners gives boosting more flexibility than other shrinkage estimators or model-selection techniques. As an extreme example, if one uses Least-Squares base-learners, *b*_OLS_, and runs the boosting algorithm until *m → ∞*, this unpenalized model will produce regression coefficients that are nearly identical to a frequentist GLM.

Practically, we deliberately constrain the base-learners and keep them weak (Bühlmann & Yu, 2003). Base-learners need only have a predictive performance of slightly better than random chance for the entire ensemble to be strong (Schapire, 1990; Kearns & Valiant, 1994). The boosted ensemble results in a smooth additive model of adaptive complexity:

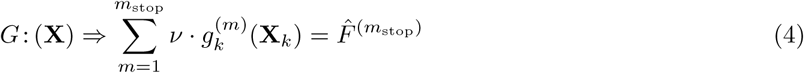

where each prediction function *g*_*k*_ is deliberately shrunk by the scalar parameter *ν ∈* (0, 1), called the *learning-rate*.

##### Conventional boosting

There are many flavours of boosting, but they all share a basic algorithm. The goals are: i) to estimate the fit-vector 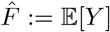, which is the vector of our expected values of *y*; and ii) to make an ensemble of base-learners *𝒢* that can make predictions from new covariate data. Boosting is summarized as: *i*) set the initial values of fit-vectors *F* ^(0)^ to the MLEs of the simplest model (such as the intercept-only model); *ii*) increment *m*; *iii*) use the current fit-vector 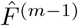 to estimate the negative-gradient of the loss-function, 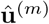 (like the residual variation unexplained by the previous step); *iv*) make a prediction function that maps *X* to 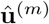 and append the prediction function to the ensemble *𝒢*^(*m*)^ *← g*\*; *v*) increment the fit-vector with the predictions from *g*\*, shrunken by the scalar *ν* such that 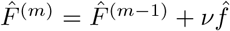; *vi*) repeat steps *ii* to *v* until *m* = *m*_stop_. The regularization parameters *m*_stop_ and *ν* govern the amount of shrinkage (Bühlmann & Yu, 2003; Schmid & Hothorn, 2008a).

##### Component-wise boosting

The development of boosting from a classification algorithm into a statistical modelling framework is credited to Bühlmann & Yu (2003). In their component-wise boosting framework, the user specifies a large candidate set of base-learners, each representing a plausible set of sub-models for different main effects and interactions and non-linear effects, etc. This is somewhat analogous to the way in which a user would set-up a large candidate set of fixed-effect models for model-selection (but simpler). Figure 3 shows a comparison of 64 different fixed-effect CJS models in Program Mark, and their equivalent representations as base-learners for CJSboost. Variable selection is integrated internally to the descent algorithm by selecting only one best-fitting base-learner per *m* iteration. In other words, base-learners compete with each other to enter the ensemble, per *m*.

**Figure 3:**
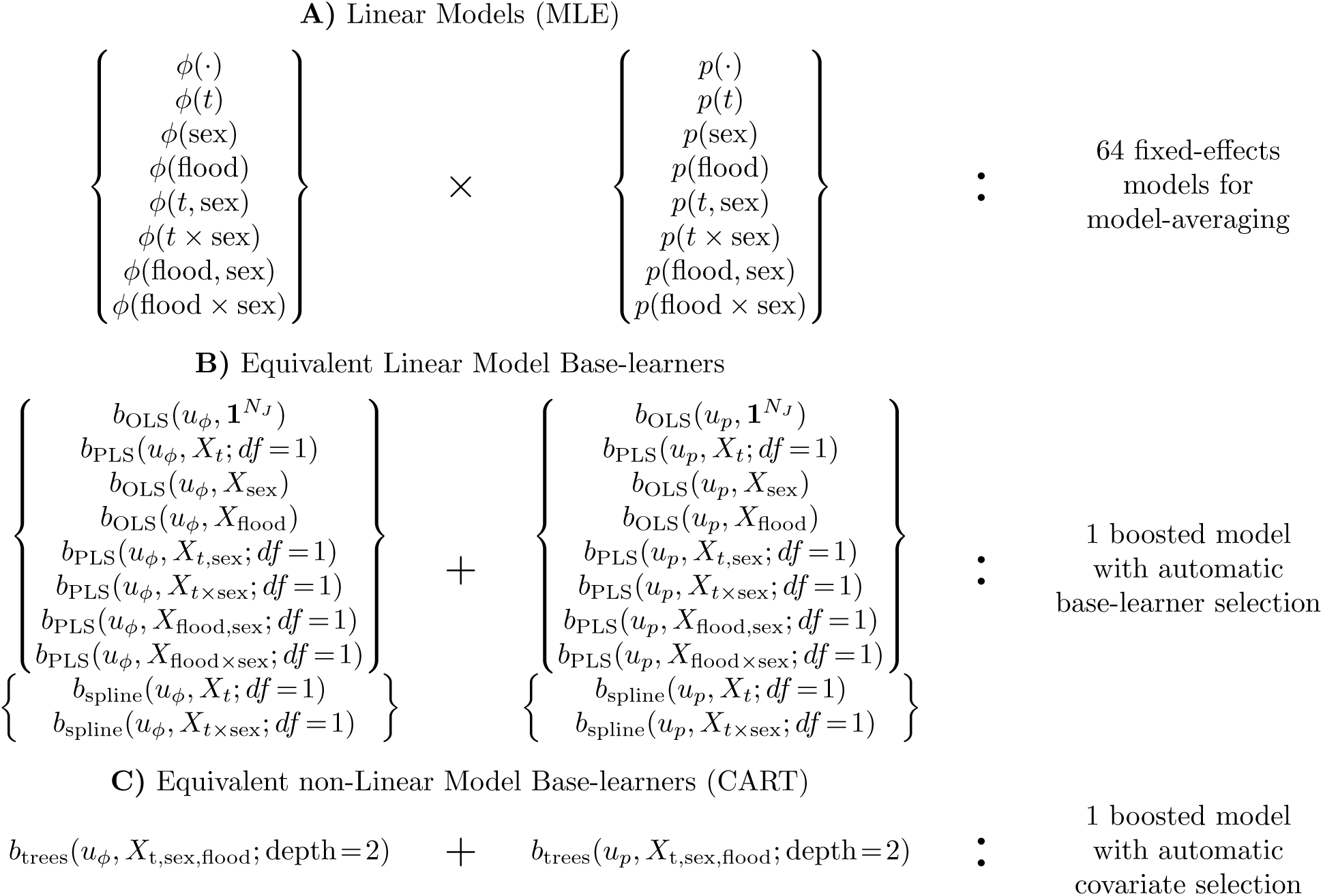
Different notation for multimodel inference of a Cormack-Jolly-Seber model, comparing fixed-effects model-averaging and boosting. **A)** Each fixed-effect model includes one term for *ϕ* (*left*) and one for *p* (*right*). *θ*(·) is an intercept model; *θ*(*t*) has different coefficients per *T* capture periods (with appropriate constraints on *t* = *T*); *θ*(*a, b*) is a linear combination of covariate *a* and *b* on the logit scale; *θ*(*a × b*) are main-effects plus the interaction between *a* and *b* on the logit scale. **B)** Equivalent linear base-learners (Ordinary and Penalized Least Squares from `mboost`; Bühlmann & Hothorn, 2007) with penalties to constrain their effective-*df*. All base-learners are available in one model; selection of base-learners is by component-wise boosting. **C)** A CJS model with CART-like trees, allowing non-linear effects and complex interactions. Selection of covariates is internal to the base-learners’ ctree algorithm (Hothorn et al., 2006).

In component-wise boosting, the fitted ensemble *𝒢* contains the final selected base-learners, which can be used to understand the functional relationships between covariate data and the response variable. For example, if covariate *x*_1_ has more predictive power than *x*_2_, we expect that the base-leaner *b*(*x*_1_) to be selected with greater frequency than *b*(*x*_2_). For least-square base-learners, we can retrieve the regression coefficient of *x*_1_ by adding up all the pertinent coefficients contained in *𝒢*, multiplied by *ν*. These have the same meaning as the regression coefficients in a GLM (except they have shrinkage). More specifically, they are almost equivalent to the regression coefficients of an *𝓁*_1_-regularizer like the Lasso (Bühlmann & Yu, 2003; Efron et al., 2004).

##### Multi-parameter boosting, or GAMLSS

Another key development was the extension of boosting to include multi-parameter likelihood functions (Schmid & Hothorn, 2008b; Schmid et al., 2010; Mayr et al., 2012), sometimes called boosted-GAMLSS (or “GAMs for location, scale and shape”). This is a wide class of interesting regression models such as Beta regression (Schmid et al., 2013) or Occupancy-Detection models (Hutchinson et al., 2011) which have multiple parameters.

The multi-parameter problem is obvious in the CJS likelihood, where we have a parameter *ϕ* for survival and a second parameter *p* for capture-probability. We must perform model-selection on both parameters. The fit-vectors **F** are no longer the expected values of the response variable *Y* (which does not interest us in CMR); instead the fit-vectors 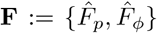 represent the expected values of the processes *ϕ* and *p* on the logit scale, 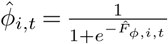. Also, we now have different ensembles of base-learners for each parameter ***𝒢***:= {*𝒢*_*p*_*, 𝒢*_*ϕ*_}.

The boosted-GAMLSS algorithm requires independent data-points, so it is not suitable for CMR. But, it provides the mechanism to jointly boost the survival and capture processes. The key innovation of boosted-GAMLSS was to estimate the negative gradient of the loss function by taking the partial derivatives of the loss function with respect to each parameters’ fit-vector, 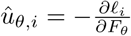, conditional on the values of the other fit vectors *F*_*θ*_.

### 2.2. CJSboost

CJSboost combines all the aforementioned ideas of conventional boosting (functional gradient descent by taking small regularized steps) and component-wise boosting (integrated variable selection) and multi-parameter boosting (interweaving boosting steps for *ϕ* and *p*), but requires one more step to make boosting applicable to CMR data. We must break the serial-dependence among individual captures within a capture-history. In other words, we garner conditional independence of data-points, and then proceed with gradient descent.

I developed two algorithms to achieve this conditional independence. CJSboost-MC uses stochastic imputation of latent states; it is described in Appendix A. I will focus on another algorithm, CJSboost-EM, which imputes and iteratively updates the expected values of latent states through an Expectation-Maximization step.

#### 2.2.1. The Expectation-Maximization Step

Expectation-Maximization (EM) is a common technique to adapt boosting to complex non-linear models, such as Conditional Random Fields (Dietterich et al., 2004), presence-only species distribution models (see the Appendix of Ward et al., 2009), and Generalized Additive Mixed-Models (Groll & Tutz, 2012).

The motivation is thus: our loss function, the negative CJS log-likelihood (1), can only be evaluated *per capture-history*, and not per data-point/capture. Therefore, it cannot be boosted because there is no point-wise evaluation of the negative gradient. As a technical remedy, we use a slightly different *surrogate* loss function which can be evaluated per data-point. This surrogate loss function is derived from the negative *Complete-Data log-Likelihood* (CDL). The CDL can be evaluated per capture because it assumes that we know the latent states (*z*_*i,t*_*, z*_*i,t*–1_) at *t* and *t*–1. The negative CDL is:

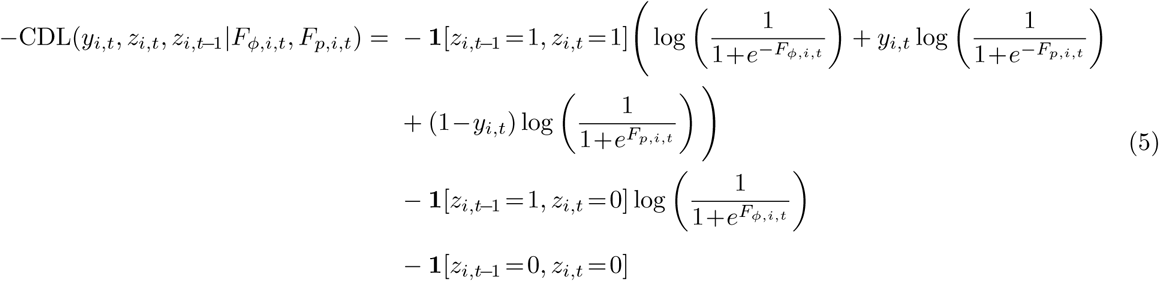

where *y* and *z* are defined as above in (1) and 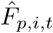 and 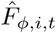 are the fit-vectors for the capture-probability and survival parameters, respectively, on the logit scale.

Using the negative CDL, we derive the surrogate loss function for the EM-step. It is called a “Q-function”. The idea is to replace the values of (*z*_*i,t*–1_*, z*_*i,t*_) in (5) with their *two-slice marginal* expectations: *w*_*t*_(*q, r*):= *p (z*_*t*–1_ = *q, z*_*t*_ = *r|***y**, **F**). *w*_*t*_(*q, r*) is the joint probability of *z*_*t*–1_ = *q* and *z*_*t*_ = *r*, conditional on the fit vectors **F** and the data **y**. The two-slice marginals {*w*(1, 1)*, w*(1, 0)*, w*(0, 0)} can easily be computed with a standard “forwards-backwards” HMM algorithm (Rabiner, 1989; Murphy, 2012b), as detailed in Appendix B. This must be done in-between boosting steps.

To simplify notation, we will index each capture *y*_*i,t*_ of individual *i* at time *t* with the index *j*:= (*i, t*). This also emphasizes how each capture is conditionally independent given *z*. The Q-function is:

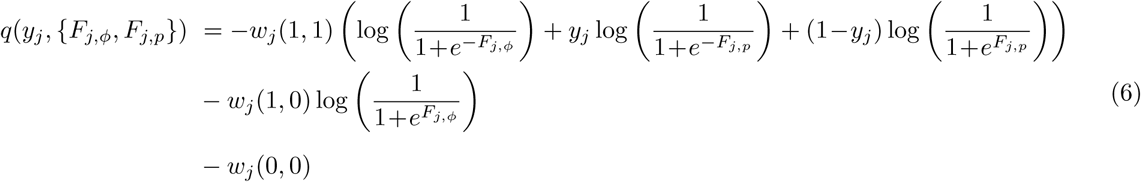

The *q* formula has a clear intuition: we are *weighting* three conditional loss functions that represent the three plausible latent state transitions: *alive → alive*, vs. *alive → dead*, vs. *dead → dead* (the fourth scenario of *dead → alive* is not permissible).

According to the theory of EM, by minimizing the surrogate loss function *q*, we also minimize the target risk function: the negative CJS log-likelihood (1). The advantage of working with the surrogate loss function is that it is easy to calculate its point-wise gradient using partial derivatives: 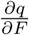(7).

The two-slice marginal expectations *w*(·, ·) change with every update of 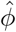 and 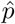. Therefore, we iteratively boost the parameters *ϕ* and *p* conditional on *w*(·, ·), and then update *w*(·, ·) conditional on 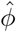 and 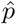. The expectations quickly converge and we fit a statistical CMR model that is optimal at prediction and has integrated variable selection.

#### 2.2.2. CJSboost-EM algorithm

The formal CJSboost-EM algorithm is as followed. It is identical to the multi-parameter component-wise boosting algorithm of Schmid et al. (2010, *§*2), except for the additional EM-step (Step 5) and, of course, different loss and gradient functions (Step 6).

1. Specify the candidate set of plausible base-learners 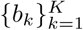, per *ϕ* and *p*.
2. Set the regularization parameters, *m*_stop_, *ν*_*ϕ*_ and *ν*_*p*_; e.g. *m*_stop_ = 10^3^; *ν*_*ϕ*_ = 0.01.
3. Initialize the fit vectors at the MLEs of a simple intercept-only model

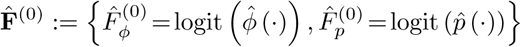
4. Set *m* = 1.
5. Estimate the two-slice marginal probabilities 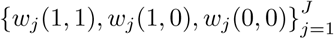 for all individuals and capture-periods, using the forwards-backwards algorithm (see Appendix B.3).
6. Estimate the gradients of the surrogate loss function *q* w.r.t the fit vectors 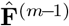:

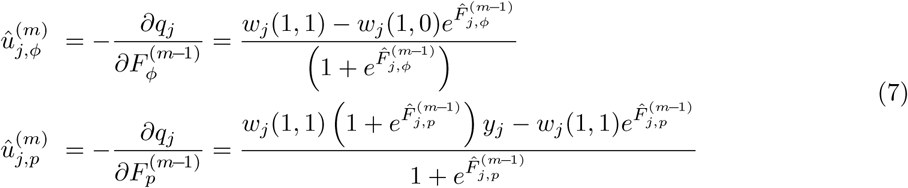
7. For each parameter *θ* in {*ϕ, p*}, do:

a. for each *k* base-learner for *θ*, do:

i. fit the base-learner to the gradient: 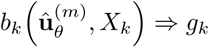;
ii. make an estimate of the gradient, 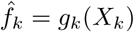;
b. find the base-learner that best-fits the gradient:

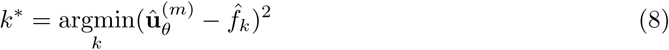
c. append the prediction function of *k*\* to the ensemble 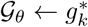;
d. re-estimate the fit vector: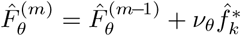;
8. Monitor the empirical risk on the full data 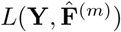. Or, monitor the holdout-risk using an out-of-sample subset of the data 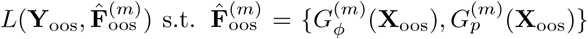 to use for bootstrap-validation.
9. Update *m* = *m* + 1.
10. Repeat steps 5 to 9 until *m* = *m*_*stop*_.

The three regularization parameters *m*_stop_, *ν*_*ϕ*_, *ν*_*p*_ control the shrinkage, and must be tuned by minimizing the average holdout-risk. This is our estimate of the expected loss (see 2.2.3).

The outputs of the algorithm are the fit vectors 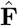 and the ensemble of fitted base-learners *𝒢* _*ϕ*_ and *𝒢* _*p*_. We can estimate the survival of individual *i* at time *t* by back-transforming the fit-vectors onto the probability scale: 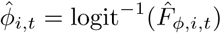. We do the same for capture-probability 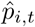. For abundance, we use the Horvitz-Thompson-type estimator: 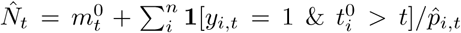 (McDonald & Amstrup, 2001). For predicting *ϕ*\* and *p*\* on new covariate data **X**\*, we merely process the data through the ensemble of fitted base-learners and shrink by *ν*, i.e., 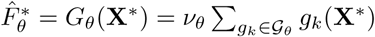.

The second algorithm, CJSboost-MC, is described in Appendix A.

#### 2.2.3. Regularization parameters

In multi-parameter boosting, the most important regularization parameters are *m*_stop_, *ν*_*ϕ*_, *ν*_*p*_, which control the shrinkage. To guarantee a prediction optimal model, we must tune *m*_stop_, *ν*_*ϕ*_, *ν*_*p*_ with cross-validation or bootstrap-validation. As per Schmid et al. (2013), I suggest bootstrapping the individual capture-histories between 50 to 100 times, training a new model on each bootstrap sample. On average, each bootstrap leaves 36.5% of the capture-histories unused in the model fitting, which can then be used to estimate a holdout-risk.

Finding the optimal value of *m*_stop_ is straight-forward and routine in conventional boosting. See Figure 1 for an example of bootstrap-validation used to estimate *m*_cv_. Tuning the Real-valued *ν*_*p*_ and *ν*_*ϕ*_ is computationally expensive and requires some careful consideration. This challenge is inherent to all multi-parameter boosting algorithms, including boosted-GAMLSS models (Schmid et al., 2013; Mayr et al., 2012) and CJSboost. Practitioners should see Appendix C for my proposed method and other ideas.

Finally, there are complexity parameters associated with individual base-learners that must be decided *a priori* and could be considered as regularization parameters, e.g., the effective-degrees-of-freedom of a Penalized Least-Squares base-learner, or the maximum tree-depth of a conditional inference tree. The effects of these parameters have been studied in conventional component-wise boosting (Bühlmann & Yu, 2003; Schmid & Hothorn, 2008a; Kneib et al., 2009). Practitioners should read Appendix D for best-practises, as well as the tutorial by Hofner et al. (2012).

### 2.3 Sparsity and Consistency

The previous discussions were predicated on prediction and minimizing the expected loss of estimation. There is another type of *model-identification* inference which is focused on finding the “correct” model, such as declaring one covariate to be truly influential and another covariate to be spurious (what Aho et al., 2014 calls “B-type” thinking). If we declare a spurious covariate to be important, it is a False Discovery (FD). If we declare a truly influential covariate to be unimportant, it is a False Rejection (FR). For this type of inference, the chief desirable property is *consistency*: that we will, with high probability, find the correct model as sample size increases. It turns out that prediction-optimal methods are generally not consistent estimators and will tend to produce FDs (Shibata, 1980, 1986a; Shao, 1993, 1997; Yang, 2005; van Erven et al., 2012)

Whether or not a procedure is consistent and/or efficient is mediated by one’s assumptions about the dimensionality of the true generative process (i.e., the number of parameters in the true model). Consistent procedures assume *sparsity*: the true generative model has a finite number of covariates, most covariates have zero effect, and the dimensionality stays constant as sample size increases. The truth is the truth regardless of sample size. This is a controversial assumption (Burnham & Anderson, 2004). For example, some authors believe that the truth is never sparse: natural phenomena are complex with an infinite number of influences. CMR theorists generally believe that as sample size increases, an MMI procedure should reveal more of these small influences (Otis et al., 1978, pages 50-51). The AIC and prediction-optimal methods may be consistent under this latter assumption, so long as one’s models are also approximately infinite-dimensional (Shibata, 1980). Conversely, consistent procedures, such as the BIC (Schwarz, 1978), Bayes Factors, the adaptive lasso (Zou, 2006), and twin-boosting (Bühlmann & Hothorn, 2010), will severely under-fit if the truth is not sparse (Sun et al., 2013) and will have unbounded maximum expected loss (Shibata, 1986b; Leeb & Pötscher, 2008). These distinctions have been more-or-less ignored in the ecological literature (but see Burnham & Anderson, 2004; Link & Barker, 2006; Aho et al., 2014; Galipaud et al., 2014). In the CMR field, consistency and model-identification has been much less important than estimating abundance, but some examples do exist (e.g. Pérez-Jorge et al., 2016; Taylor et al., 2016).

Interestingly, two recent papers by Meinshausen & Bühlmann (2010) and Bach (2008) have proposed similar ways to use *𝓁*_1_-regularizers, like the Lasso and boosting, in order to find truly influential covariates under “high-dimensional” situations: when the sample size is small and there are many potential covariates. The idea is to subsample/resample the data, and tally the frequency that each covariate is selected by an *𝓁*_1_-regularizer, over the entire space of the regularization parameter (e.g., *m* in boosting). Some authors have suggested that these are Frequentist approximations to Bayesian posterior inclusion probabilities (Richardson, 2010; Draper, 2010; Murphy, 2012c). I will loosely refer to these procedures as “stability selection”, although there is a lot of subtle variation in this rapidly evolving field of research. In particular, its application in multi-parameter boosting, like boosted-GAMLSS or CJSboost, is still unvalidated. See Appendix F for clarifications.

There are two key points. First, this type of MMI is no longer about prediction nor estimation, but uses prediction-optimal methods as an intermediate step for correct model-identification, i.e., which covariates are part of the true model. Second, posterior inclusion probabilities lead to straight-forward inferences: covariates with high inclusion probabilities are probably more important; covariates with low inclusion probabilities are probably not that important.

Thus, CJSboost offers a choice to CMR practitioners. If one’s goals are to estimate abundance or survival, then one can use the vanilla CJSboost model tuned for optimal prediction. Or, if one’s goals are to find covariates that significantly effect survival, then one can use the stability-selection-enhanced CJSboost and calculate inclusion probabilities. This choice is analogous to switching from the AIC to the BIC.

### 2.4 Simulation 1: Estimation

The first simulation investigated the ability of CJSboost to estimate abundance and survival, over different sample sizes. Technically, I demonstrate that minimizing the average holdout-risk also minimizes the square-error of estimating abundance and survival, as benchmarked against AICc model-selection and AICc model-averaging. I used the AICc because it is supposed to excel at precisely this kind of task: minimizing estimation error. I focused on metrics of *relative efficiency*, because this exemplifies the choice faced by Frequentist practitioners: to choose among procedures based on their relative performance to get as close as possible to the truth, over all theoretical data-sets.

I tested two CJSboost-EM models: i) a linear-model called *b*_PLS_-CJSboost, which used least-square base-learners, as listed in figure 3; and ii) a non-linear model, called *b*_trees_-CJSboost, which used conditional inference trees (Hothorn et al., 2006). The AICc-methods used 64 fixed-effects models listed in figure 3.

The simulated data-sets were inspired by the European Dipper dataset from Lebreton et al. (1992). There were *T* = 10 primary periods and two sexes of individuals (*X ∈* {1, 2}). Individuals’ first-capture periods 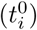 were random. The true processes were time-varying effects plus an individual sex effect (*X*). The true data-generating processes ^2^ were: *ϕ*(*t, X*) = 0.91 – 0.01 *t*– 0.05 · **1**[*t* = 5, 6] + 0.05 · **1**[*t* = 9, 10] – 0.05 · **1**[*X* = 1] and 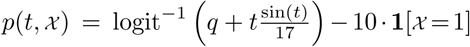, where *q* controlled the mean capture-probability. Figure 5 graphs an example simulation. For analyses, there was an additional categorical variable, called *Flood*, which grouped the captures periods {4, 5, 6}: it simulates an analyst’s hypothesis that dipper survival and capture-probability are different in periods 4, 5 and 6, due to environmental degradation by flooding.

**Figure 4:**
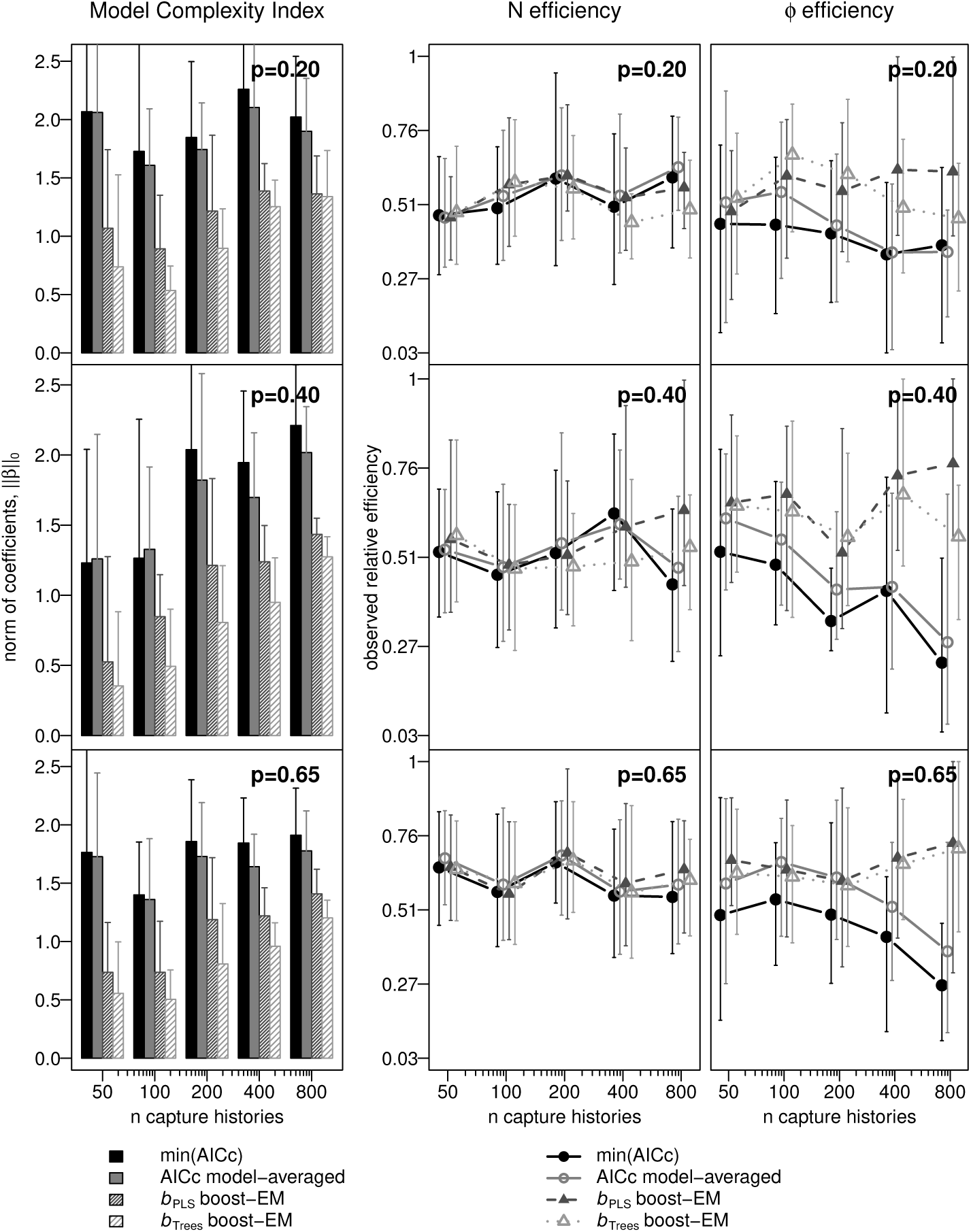
Simulations of Cormack-Jolly-Seber data-sets show how model complexity and estimation performance vary by sample-size (*x-axes*), true capture-probability (*p* = 0.2, 0.4, 0.6, *panel-rows*), and the multi-model inference paradigm: AICc methods (*thick-lines*) vs CJSboost methods (*dashed-lines*). *Left*: model-complexity increases as the sample-size increases, as measured by the absolute size of the estimated model coefficients (a.k.a the norm of 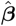). *Middle*: relative performance estimating abundance *N*_*t*_, as measured by the average *observed efficiency* MSE_min_*/*MSE *∈* (0, 1], where MSE_min_ is the error of the best estimator. Higher efficiency is better. *Right*: The average observed efficiency of survival. Results are averaged over 20 simulations per combination of *p* (*panel-rows*) and *n* (*x-axes*).

**Figure 5:**
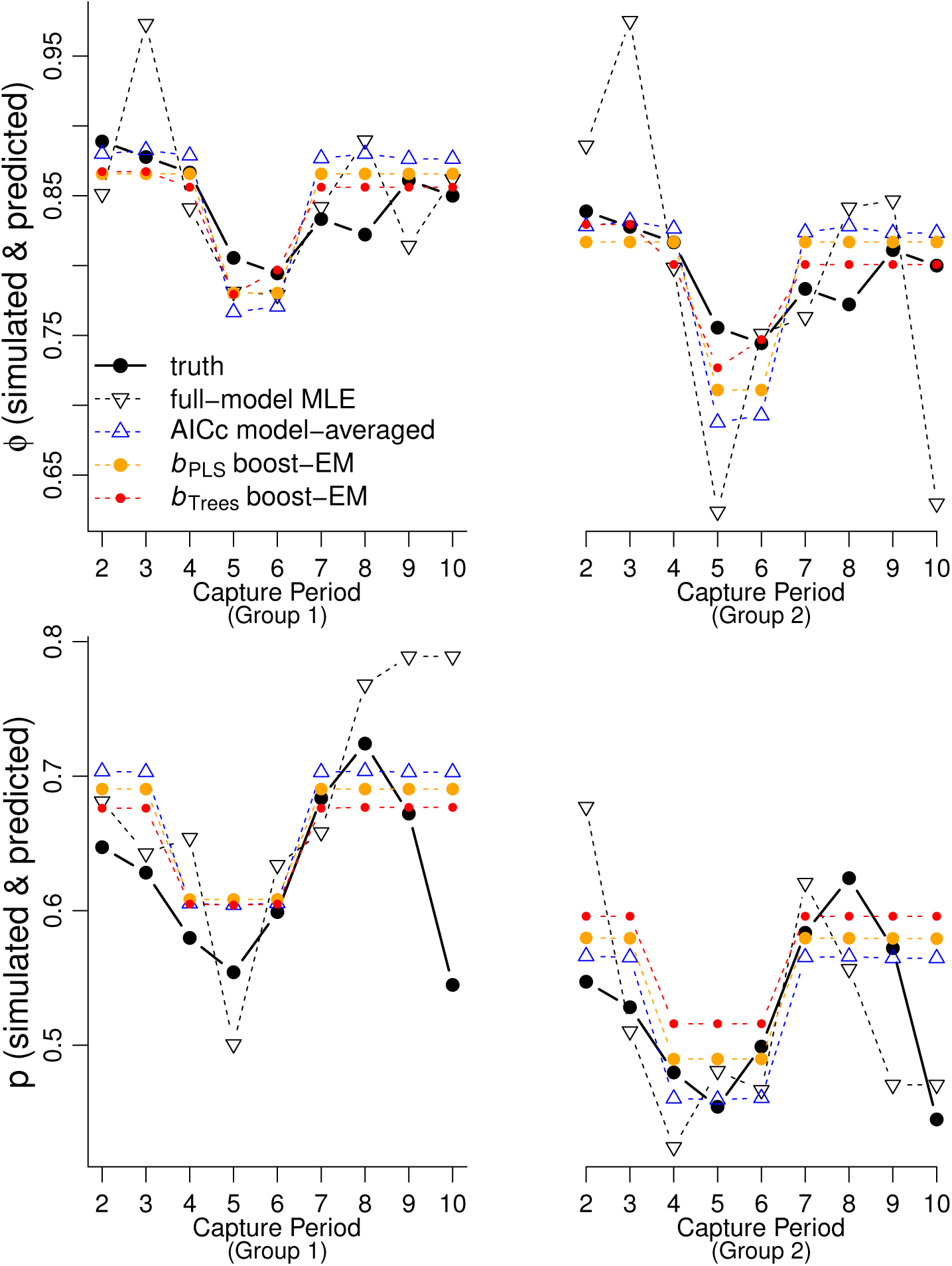
Simulation 1, demonstrating the CJSboost estimates from the Expectation-Maximization technique. A comparison of capture-probability estimates 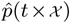 and survival estimates 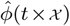 from models composed of linear base-learners (OLS and PLS; in orange) and non-linear base-learners (CART-like trees; in red), as well AICc model-averaging (blue) and MLE (dashed black).

For each simulation and estimator, the mean standardized square error (MSE) was calculated for abundance (*N* _*t, x*_) and survival (*ϕ*_*t, x*_), e.g. 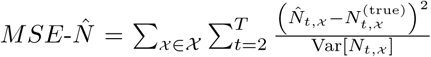. A lower MSE is better. We compared the estimators’ MSE values by two statistics: i) the *observed efficiency* of estimator *i*, which is 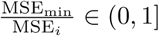 (higher is better), where MSE_min_ is the MSE of the best performing estimator; and ii) *rank*, which is the rank-order of estimates by increasing values of MSE (rank 1 is best). These criteria were used by early researchers of the AIC and BIC (Shibata, 1980; McQuarrie, 1999). Both criteria are empirical ways of approximating the more fundamental Frequentist value of relative *efficiency*. Better values imply that a procedure has, over repeated sampling, estimates that are closer to the truth (but not necessarily unbiased).

The observed efficiency and rank calculations were summarized according to sample size scenarios: different combinations of average capture-probabilities *p ∈* {0.2, 0.4, 0.65} and the number of captured individuals *n ∈* {50, 100, 200, 400, 800}. I ran 20 simulations per combination of *n* and *p*.

All boosting models used 70-times bootstrap-validation to estimate optimal values of *m*_stop_, *ν*_*ϕ*_ and *ν*_*p*_. The base-learners were taken from the mboost R package (Bühlmann & Hothorn, 2007; Hofner et al., 2012). The AICc model-averaging analyses were conducted in Program MARK (White & Burnham, 1999) and RMark (Laake, 2013).

### 2.5. Analysis: Dipper Example

Using CJSboost-EM, I reanalyzed the European Dipper dataset from Lebreton et al. (1992). I compared the results to the MLEs of the fully-saturated model (*ϕ*(*t ×* sex)*p*(*t ×* sex)) as well as to AICc model-averaged estimates. The dataset has 294 individuals in *T* = 7 capture periods. Covariates included time, sex, and flood, similar to Section 2.4. The model-building framework was the same as in Figure 3. 100-fold bootstrap-validation was used to optimize *m*_stop_, *ν*_*ϕ*_ and *ν*_*p*_.

Interested readers can repeat this analysis using the online tutorial at http://github.com/faraway1nspace/HMMboost/.

### 2.6 Simulation 2: Sparsity and Consistency

The final simulation addressed the issue of high-dimensionality and the ability of CJSboost (EM) to find a sparse set of important covariates out of many spurious covariates. This type of *model-identification* inference is distinct from the estimation/prediction goals of shrinkage estimators and AIC approaches. The loss-function is no longer about minimizing a square-estimation error, but is focused on limiting False Discoveries (FD) and False Rejections (FR). For this task, one desires an estimator that is *model-selection consistent* ; which is to say, it will make zero FDs and FRs with probability 1 as sample size gets large.

Practically, this challenge is inappropriate for fixed-effect model-selection, because one must consider all combinations of covariates for different parameters (*ϕ, p*). In this section, I simulated 21 multi-collinear covariates, resulting in more than 4 trillion different fixed-effects models (excluding two-way interactions). It is clearly impossible for all-subsets model-selection (unless one takes ill-advised short-cuts).

#### 2.6.1. Stability Selection and Inclusion Probabilities

Theoretically, this challenge is also inappropriate for the vanilla CJSboost or other shrinkage estimators. Instead, I propose to use a bootstrapped-enhanced CJSboost to produce a consistent estimator. The crux of this estimator is to approximate the Bayesian probability that a covariate is part of the “true model”, a.k.a. posterior inclusion probabilities, *π*(*I*_*θ,k*_*|***Y**, **X**). We desire such probabilities because they lead to inferences about the significance of covariates. ^3^ Influential covariates should have very high inclusion probabilities that converge to 1 at high sample sizes, while spurious covariates should have low probabilities. In this simulation, I will show the distribution of inclusion probabilities for truly-influential and spurious covariates, as a function of different sample sizes.

Inclusion probabilities are a fundamentally Bayesian quantity, but Frequentist approximations are desirable for significance testing in a multi-model framework (Lee & Boone, 2011). Some authors (Richardson, 2010; Draper, 2010; Murphy, 2012c) noticed that such an approximation is possible through Stability Selection plus *𝓁*_1_-regularization (Meinshausen & Bühlmann, 2010; Shah & Samworth, 2013). The idea is to subsample/resample the data and tally the number of times that a covariate is selected by an *𝓁*_1_-regularizer, over all values of the regularization parameter (*m*, *ν*_*ϕ*_, *ν*_*p*_). To calculate the approximate inclusions probabilities, 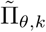, I propose the following procedure: set the values of *ν*_*ϕ*_ and *ν*_*p*_ to their prediction-optimal values 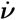; bootstrap the capture-histories *B* times; for each *b* bootstrap, run CJSboost for *m*_stop_ iterations, where *m*_stop_ *≫ m*_cv_. Stability selection probabilities, 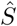, are estimated by scoring whether a *k*^th^ covariate is selected in a *b* bootstrap before *m* iterations (conditional on 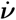), 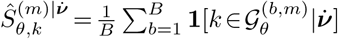. Notice that 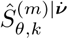 is evaluated per *m* and per covariate *k* and per parameter *θ* ∈ {ø, *p*} 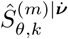 will always increase with *m* (i.e., weaker *𝓁*_1_-regularization will always increase the chance of selecting a covariate; see Figure 8). Call 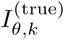 the indicator of whether the *k*^th^ covariate is part of the true model, then the inclusion probability is approximated by 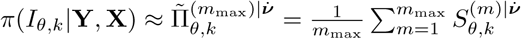.

**Figure 6:**
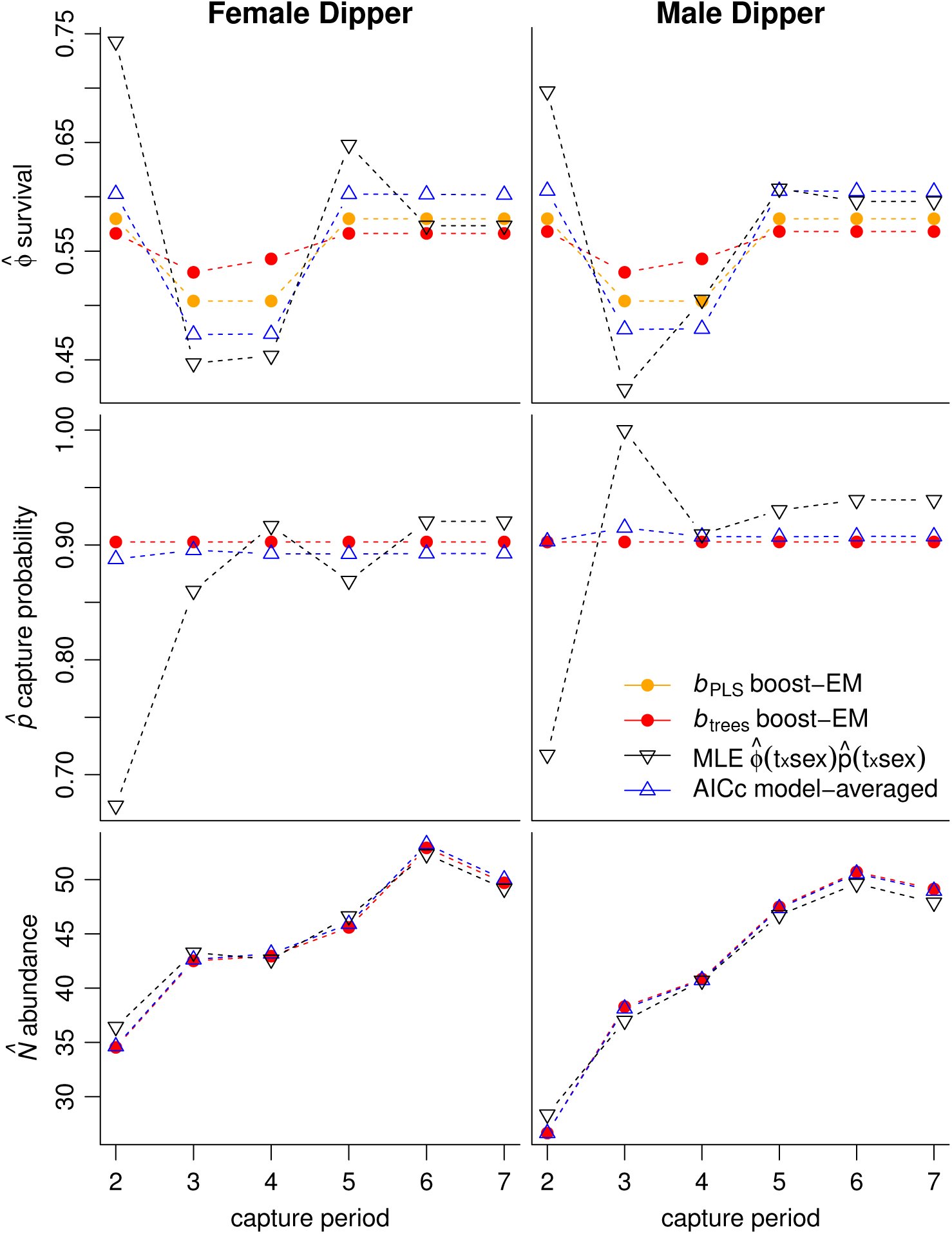
Comparison of Dipper survival 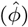, capture-probability 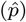, and abundance estimates 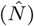 according to three predictive models: i) CJSboost-EM using penalized least-squares base-learners (PLS), ii) CJSboost-EM using non-linear conditional inference trees, and iii) AICc model-averaging in Program MARK. Plus, the MLEs of the full-model 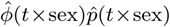.

**Figure 7:**
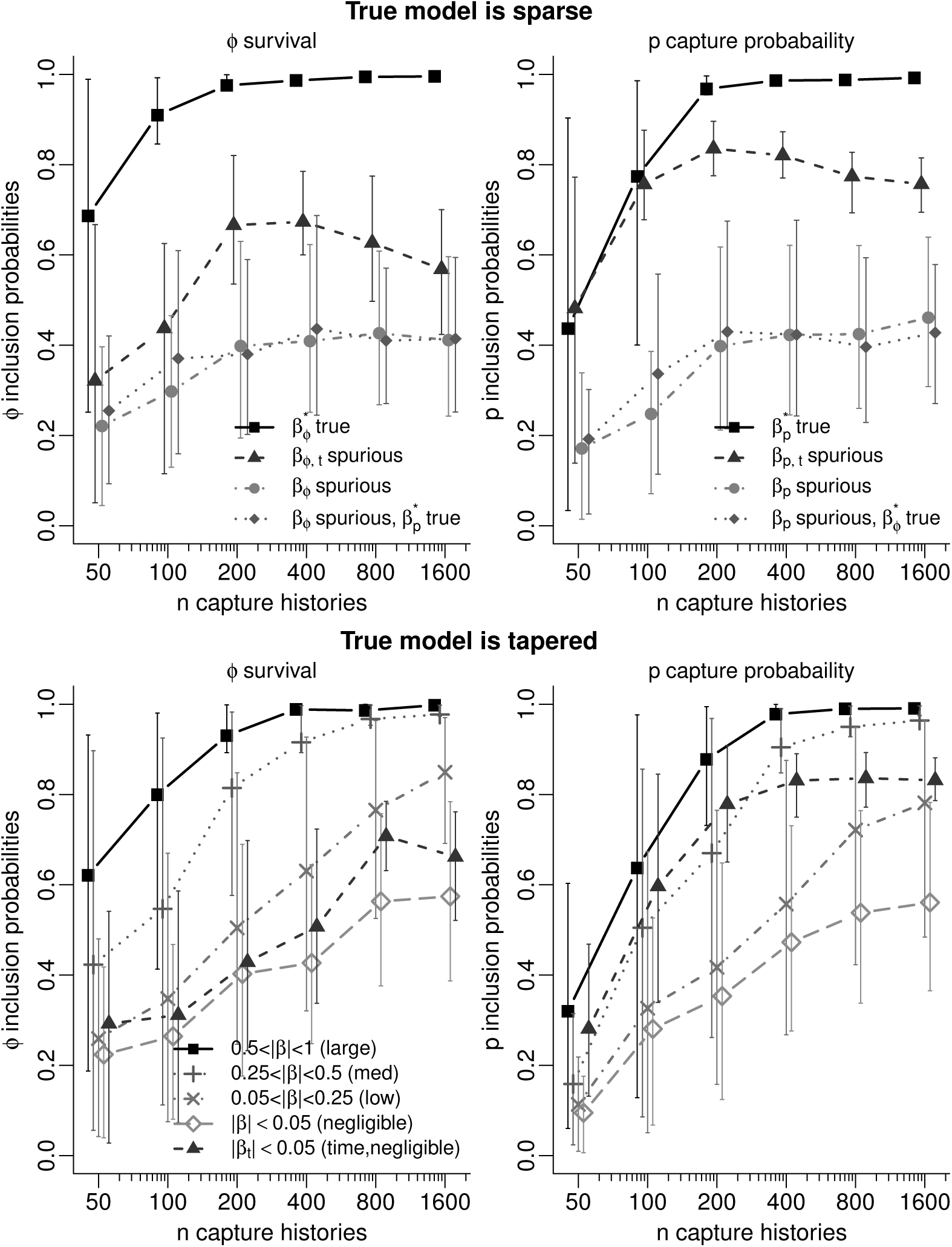
Results of 240 simulations to demonstrate the usefulness of approximate inclusion probabilities (on *y-axes*) for inference about which covariates are truly influential (i.e. part of the true model) vs. spurious covariates, over different sample sizes (*xaxes*). Each dot is an average inclusion probability over 20 simulations. Scenario A (*top*): the true model is sparse: only three covariates out of 22 are truly influential on *ϕ* or *p* (*black squares*); others are spurious (*grey circles*); some are spurious for *ϕ* but influential on *p* (*grey diamonds*) and vice-versa. Time-as-a-categorical variable, when spurious, is also plotted (*dark triangles*). Scenario B (*bottom*): the true model is tapered: all 22 covariates have some contribution to the *ϕ*/*p*-process, but they vary in the magnitude of their marginal effects (*|β*_*k*_*|*). Bars are *≈ ±*1S.D.

**Figure 8:**
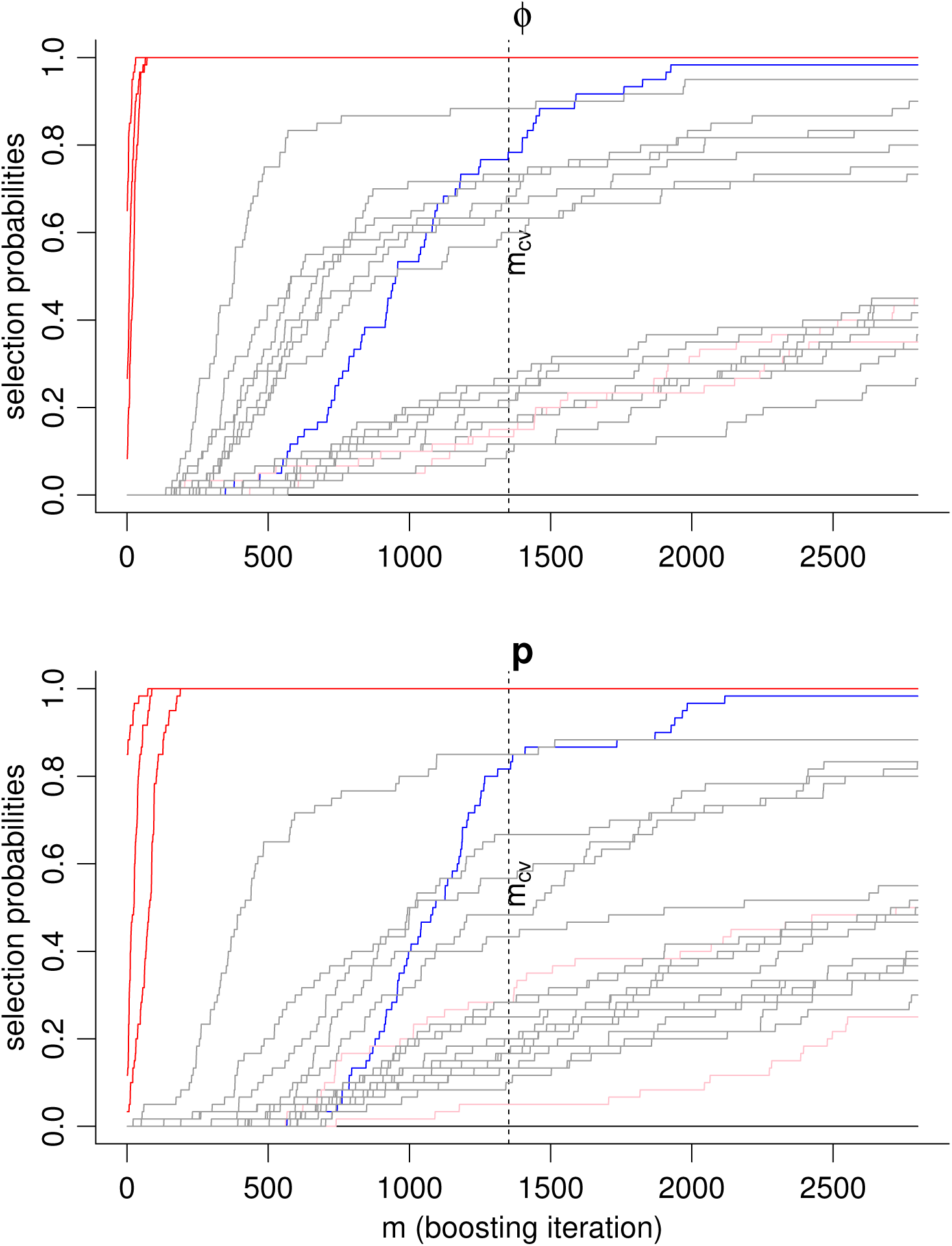
Demonstration of stability selection probabilities for one high-dimensional simulation. As the boosting iteration (*m*) gets large, regularization gets weaker, and all covariates have a higher selection probability *S* (estimated from a bootstrap). Lines in **red** are truly influential covariates. Lines in **gray** are non-influential covariates. Lines in **pink** are not-influential for *θ*, but are influential in the other parameter *θ*. Lines in **blue** represent the time-as-a-categorical-variable base-learner, a.k.a *θ*(*t*), which in this simulation was non-influential.

From a Bayesian perspective, it is like we have a prior distribution on the model-coefficients that is the exponential of the negative regularization parameter (*m*) (Geman et al., 1992), and we are crudely integrating over the prior to score selection indicators. Technically, we should integrate over *ν*_*ϕ*_ and *ν*_*p*_ as well as *m*. I propose focusing on *m* strictly for computational convenience, but this short-cut needs further validation. Readers should refer to Appendix F to see how the above formulation relates to the existing literature on stability selection (Bach, 2008; Meinshausen & Bühlmann, 2010; Schmid et al., 2012; Shah & Samworth, 2013; Hofner et al., 2015).

#### 2.6.2. Simulating Data

In 240 simulations, I use the following generative model for survival and capture-probability:

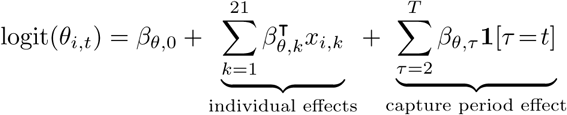

The intercepts were drawn randomly from *β*_*p,*0_ *∼* U(0.4, 0.6) and *β*_*ϕ,*0_ *∼* U(0.55, 0.8). I simulated 21 multicollinear covariates (18 continuous, three discretized) drawn from a multivariate Gaussian with marginal variances of 1 and off-diagonal correlations between 0 to 0.6. Time-as-a-categorical-variable 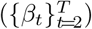 was also included as a possible influential covariate, for a total of 22 “covariates”. The number of captured individuals was stratified as *n ∈* {50, 100, 200, 400, 800, 1600}. There were *T* = 10 capture periods.

The values of the true ***β*** coefficients were drawn randomly according to two different scenarios: A) *sparsity*, in which case a few ***β***\* values were large but most ***β*** values were zero (i.e., many spurious covariates); and B) *tapering*, in which case the values of the *|****β***\**|* decreased exponentially from one or two large values, to many small-but-nonzero values. I ran 120 simulations per scenario A and B. I highlight these scenarios because sparsity is a fundamental assumption of all model-selection consistent procedures, whereas some authors suggest that tapering is more in-line with reality (Burnham & Anderson, 2004). Tapering also challenges the very notion of a “true model”, in which case we can only speak about the best approximating model (but see Link & Barker, 2006). In an extreme form of tapering, when the magnitudes of the *β* values actually increase with sample-size, consistent procedures can have a worst-case estimation error that becomes infinite (Leeb & Pötscher, 2008), which I highlight to remind practitioners of the price of this type of multimodel inference.

For the sparsity scenario (A), three covariates were randomly picked to have a significant effect, i.e. 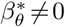. These truly influential covariates, 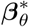, had norms of 1 on the logit scale, resulting in large marginal effects 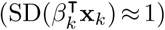 that spanned 0.8–0.9 probability-units. When the 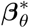 were categorical variables, then they had norms of 3 in order to achieve a similar marginal effect. The coefficients were simulated separately for *ϕ* and *p*.

**Table 1:**
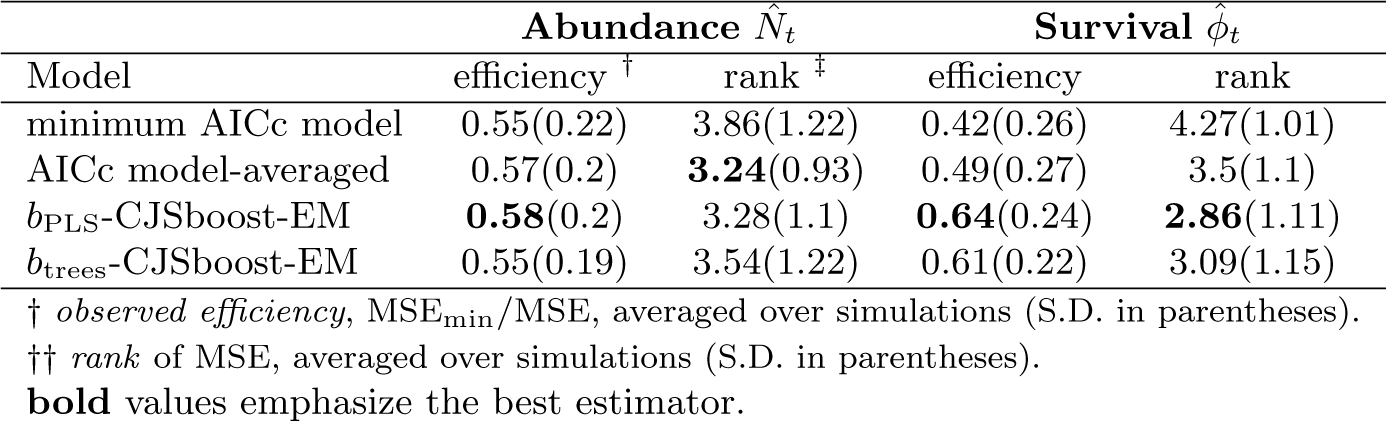
CJSboost vs AICc for estimating survival and abundance: results of simulation 1.

| Model | Abundance $\hat{N}_t$ | | Survival $\hat{\phi}_t$ | |
| --- | --- | --- | --- | --- |
|  | efficiency <sup>†</sup> | rank <sup>‡</sup> | efficiency | rank |
| minimum AICc model | 0.55(0.22) | 3.86(1.22) | 0.42(0.26) | 4.27(1.01) |
| AICc model-averaged | 0.57(0.2) | **3.24**(0.93) | 0.49(0.27) | 3.5(1.1) |
| $b_{\text{PLS}}$-CJSboost-EM | **0.58**(0.2) | 3.28(1.1) | **0.64**(0.24) | **2.86**(1.11) |
| $b_{\text{trees}}$-CJSboost-EM | 0.55(0.19) | 3.54(1.22) | 0.61(0.22) | 3.09(1.15) |

<sup>†</sup> *observed efficiency*, $\text{MSE}_{\min}/\text{MSE}$ , averaged over simulations (S.D. in parentheses).
<sup>‡</sup> *rank* of MSE, averaged over simulations (S.D. in parentheses).
**bold** values emphasize the best estimator.

For the tapering scenario (B), all *β*_*θ*_ values were non-zero. On average 5.6% of ***β*** had marginal effects categorized as “large” 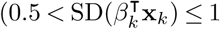, or equivalently 0.5 *<|β*_*k*_*| ≤* 1), 13.9% were “moderate” (0.25 *<|β*_*k*_*| ≤* 0.5), 37.3% were small (0.05 *< |β*_*k*_*| ≤* 0.25) and 43.1% were negligible (0 *< |β*_*k*_*| ≤* 0.05). The coefficients were simulated separately for *ϕ* and *p*.

#### 2.6.3. Data Analysis

To analyze each simulated dataset, I use the following base-learners for each *p* and *ϕ* sub-model: 22 PLS base-learners (*df* = 2) for each continuous and categorical covariate; a PLS base-learner for the time-as-a-categorical variable (a.k.a, the *θ*(*t*) model); and a base-learner for the intercept. In stability selection, base-learners must have equal flexibility/degrees-of-freedom; otherwise, the more complex base-learners will have a greater probability of being selected (see Section 2.2.3). The regularization parameters *ν*_*p*_ and *ν*_*ϕ*_ were optimized with ten 70-fold bootstrap-validation exercises, as per Section Appendix C.1.

#### 2.6.4. Oracle Estimator

Finally, an auxiliary task was to derive an *oracle estimator* (Fan & Li, 2001; Zou, 2006). The goal is estimate the coefficients as if we knew the “true” model from the beginning, a property of all consistent procedures (Leeb & Pötscher, 2008). The idea is to threshold the inclusion probabilities at some high threshold *π*_thr_ *∈* (0.5, 1), and use only those covariates where 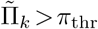 (called hard-thresholding). A final un-regularized CJSboost model is used to get “debiased” estimates by running *m → ∞* (Bach, 2008; Murphy, 2012c) ^4^. I showcase this oracle property on just one simulated dataset from scenario A, in order to demonstrate the role of the threshold *π*_thr_ in determining the oracle properties and the number of FDs and FRs.

## 3. Results

### 3.1. Simulation 1: CJSboost vs AIC

Table 1 and Figure 4 summarize the estimation performance of boosting-EM and AICc methods across all simulations. Figure 5 shows the model fits and the true processes for one example simulation (*n* = 300).

The general result is that the *b*_PLS_-CJSboost model with PLS base-learners did best at minimizing estimation errors and obtaining higher relative efficiencies for both abundance 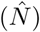 and survival 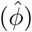, over all samples sizes, followed by AICc model-averaging, then *b*_trees_-CJSboost with conditional inference trees. The worse performance was by the minimum AICc model.

Regarding abundance estimates, all four estimators had similar performances, with no discernible trend by sample size (*n* and *p*). *b*_PLS_-CJSboost had slightly better performance according to the observed efficiency criteria, while AICc model-averaging won narrowly according to the average MSE rank.

However, for survival, the CJSboost models clearly outperformed the AICc methods, especially with the PLS base-learners: they obtained the highest overall efficiencies and best mean rank. The results varied by *n*: when *n ≤* 100, all methods had similar performances; but when *n >* 100, the boosting methods greatly out-performed both AICc methods.

To understand why boosting out-performed the AICc methods, it is helpful to look at the growth in the magnitude of the model coefficients (*‖****β****‖*). According to theory on shrinkage, we would expect that *‖****β****‖* would be smaller at low *n* and low *p*, for both boosting and AICc methods, to prevent over-fitting. The AIC methods had more extreme coefficient values, especially at low *n* and low *p*. Therefore, AIC methods were *under* estimating the correct amount of shrinkage necessary for optimal estimation. The *b*_trees_ models had slightly lower coefficient norms than the better performing PLS models, which suggests that the tree-models were *over* estimating the correct amount of shrinkage.

Interestingly, AICc model-averaging produced better estimates than the best AICc model, with more shrinkage on coefficients. This is unsurprising for estimating abundance. However, there are theoretical problems with model-averaging when it comes to estimating model parameters such as survival, especially under collinearity (Cade, 2015) which is an inherent feature of CMR processes. At low sample sizes (*n* = 50) both AICc methods had very high coefficient values, and a lot of variability. This may suggest that the AICc approximation does not hold well for CMR models at very low sample sizes. Interestingly, the abundance estimates were still competitive with boosting.

We can gain more insights into shrinkage by scrutinizing one example simulation (Figure 5). None of the estimators did a convincing job of approximating the true underlying processes. The estimates from boosting-EM and AICc-methods revealed similar patterns for both for *ϕ* and *p*, but they differed in the amount of shrinkage: the boosted estimates were *shrunk to the mean* more than model-averaged estimates. More shrinkage resulted in better MSE performance (despite the increase in bias). The tree base-learners had perhaps too much shrinkage and worse MSE. The Figure also shows the MLEs to illustrate the bias-variance trade-off: the MLEs of the full-model 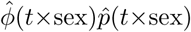 are unbiased but are also high-variance, in the sense that the estimates vary wildly around the true processes.

Figure 5 has been repeated in Appendix A using the the Monte-Carlo CJSboost algorithm.

### 3.2. Results: Dipper example

This section shows the reanalysis of the European Dipper dataset from Lebreton et al. (1992) by CJSboost-EM. Comparisons were between the linear *b*_PLS_-CJSboost model and the nonlinear *b*_Trees_-CJSboost model as well as model-averaged estimates by AICc, and the MLEs from the full-model *ϕ*(*t ×* sex)*p*(*t ×* sex). See Figure 6 for the fitted processes. The results can be summarized:

1. For both survival *ϕ* and capture-probability *p*, the three predictive methods (AICc, *b*_PLS_-CJSboost or *b*_trees_-CJSboost) had similar patterns, unlike the full-model MLE. The predictive models differed according to the amount of shrinkage.
2. The *b*_trees_-CJSboost model applied a lot shrinkage towards the time-constant values. Whereas the AICc model-averaged estimates had less shrinkage and were more similar in pattern to the MLEs of the full-model. The *b*_PLS_ model had shrinkage that was intermediate between the AICc and *b*_trees_ estimates.
3. For survival, all three predictive methods yielded the same estimates: a survival probability of 0.48-0.5 during the flood years (*t* = 3, 4) and little-to-no sex-effect (*<* 0.005 difference between male and females).
4. For capture-probability, the model-averaged estimates suggested a slight sex effect of about 1.5 probability units, whereas both boosted models shrunk the capture-probability to a constant; in contrast, the MLEs varied much more.
5. Abundance estimates showed little variation among methods, due to the high overall capture-probabilities (*p ≈* 0.9).

### 3.3. Simulation 2: sparsity, consistency, and high-dimensional data

Figure 7 summarizes the results of 240 high-dimensional simulations and their inclusion probabilities 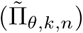 for truly influential and spurious covariates. The figure stratifies the average inclusion probabilities by sample size (*n*), parameter *θ ∈* {*ϕ, p*}, marginal effect sizes (*|β*_*θ,k*_*|*), and by the nature of the true model (*sparsity* vs *tapering*). I remind readers that we desire 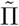 values of the truly influential covariates to converge to 1 and be well separated from the 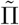 values of the spurious covariates.

The results are summarized according to the nature of the true model.

1. When the true model was *sparse* (i.e. three high-magnitude covariates and many spurious covariates) the results were:

a. For survival, there was a good separation of the 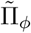 values between the truly influential covariates and the spurious covariates, when sample sizes were *n ≥* 100. Ideally, we would prefer that the *minimum* 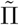 of influential covariates is high and the *maximum* 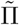 of spurious covariates is low. The average minimum 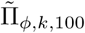 of the true covariates was 0.77 at *n* = 100, and grew to *≫* 0.9 for *n >* 200. The average maximum 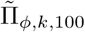 of the spurious covariates was 0.64 at *n* = 100 and grew to *≈* 0.75 at greater sample sizes. For spurious covariates, the overall average 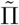 stabilized and plateaued below 0.5, while for the true covariates, the 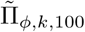 values converged to 1 for *n >* 200.
b. For the covariates influencing capture-probabilities, there was less separation of the 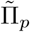 values between true covariates and spurious covariates, although the true covariates had 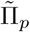 values which converged to *≈* 1 by *n >* 200, and the spurious covariates remained below 0.5.
c. The time-as-a-categorical variable (***β***_*ϕ,t*_ and ***β***_*ϕ,p*_), when spurious, had higher average 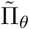 values than the other spurious covariates. For *ϕ*, the average *maximum 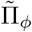* for ***β***_*ϕ,t*_ was generally between 0.6 – 0.67. For *p*, the average *maximum 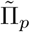* for ***β***_*p,t*_ was generally between 0.8 – 0.85. This may suggest a violation of the assumption of “exchangeability” among spurious covariates (Meinshausen & Bühlmann, 2010).
d. Covariates that were spurious in *ϕ* but truly influential upon *p* (and *vice versa*) did not seem to have 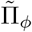 values that were different than the other spurious covariates. In other words, the true model of *ϕ* did not seem to influence the inclusion probabilities for the covariates in *p*, and *vice versa*. This suggests that the assumption of exchangeability of spurious covariates may hold in multi-parameter boosting.
2. When the true model was *tapered* (i.e. all covariates were part of the true model, but with decreasing magnitudes of marginal effects) the results were the following:

a. The overall pattern of 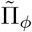 values behaved as one would expect. The covariates with *large* effects had high 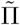 values that converged to 1 as *n* got large, while the covariates with *medium* and *small* effects had lower average 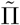 values that increased as *n* got large, and the *negligible* effects had the lowest average 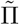 values, but which nonetheless increased as *n* got large (although their average remained below 0.5).
b. Seemingly, all effect sizes had monotonic increases in inclusion probabilities with increasing sample size. This was unlike the sparse scenario, where the 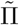 values seemed to plateau at their asymptotic distributions.
c. When time-as-a-categorical variable had negligible marginal effects, it nonetheless got higher 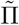 values than the other negligible covariates, especially for *p*. In other words, ***β***_*p,t*_ had a greater propensity to be selected, even when it only had a tiny marginal effect.

We can also scrutinize the results of an example simulation (sparse, *n* = 300) and visualize the stability selection pathways that were used to approximate the posterior inclusion probabilities 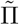. Figure (8) shows how the truly influential covariates entered the ensemble very early (small *m*) and achieved stability selection probabilities of 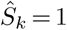. There was a lot variability in the selection pathways of the spurious covariates, but they generally increased as the amount of regularization got weaker (*m* got larger). Sometimes their 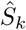 values did reach 1. Readers can view an online animated GIF which shows the stability paths for 30 example simulations, at http://github.com/faraway1nspace/HMMboost/ and in the Supplementary Material.

The point of these simulations was to show that the inclusion probabilities 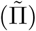 may themselves be a satisfactory end-point for an analysis. Alternatively, we can go one step further and *hard-threshold* the 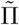 values by *π*_thr_ and discard the covariates with 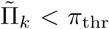. See Table 3.3. If *π*_thr_ is too low, then some spurious covariates will get selected and there are False Discoveries (FDs). If *π*_thr_ is too high, then some truly influential covariates get Falsely Rejected (FRs). Meinshausen & Bühlmann (2010) suggest that this threshold should be in the vicinity of 0.9 – 0.95, and my simulations support this threshold.

Hard-thresholding can also help us derive an oracle estimator and produce estimates that are the same as a model run with 100% foresight about the true model. This type of inference seemingly blends the two domains of MMI: estimation/prediction and consistent model-identification. Our oracle estimates are produced by: i) setting *π*_thr_; ii) discarding spurious covariates 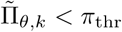; iii) and running a final CJSboost model with *m → ∞* (called “debiasing” by Murphy, 2012c, or “unregularized” by Bach, 2008). If our selection procedure is model-selection consistent, then the new estimates should have oracle properties at large sample sizes. This seems to be the case when the thresholds are high (0.8 *< π*_thr_ *<* 0.99), and both FDs and FRs are zero. However, readers should heed the warnings of Leeb & Pötscher (2008) who proved that oracle estimates can be very inaccurate at low-to-medium sample sizes, especially if the true model is not sparse. In other words, the maximum expected loss is unbounded. This is intuitive: just because we know the correct model, does not mean we can accurately estimate its true effect.

**Table 2:**
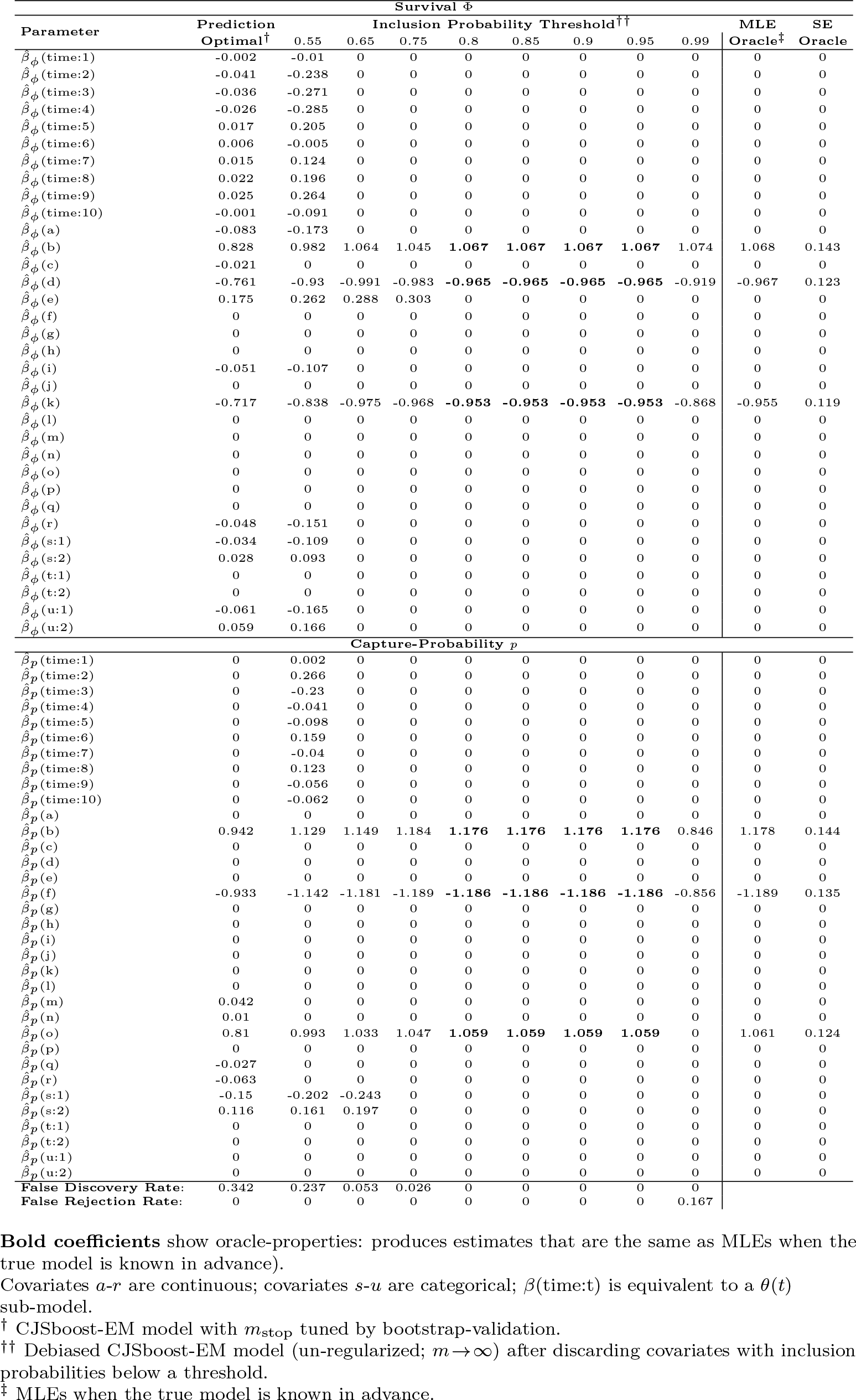
Estimates of coefficients from CJSboost, for one high-dimensional model-selection problem, under different degrees of hard-thresholding.

## 4. Discussion

This study presents CJSboost: a type of multi-model inference technique for a class of Hidden Markov Models (HMMs) known as capture-mark-recapture (CMR). I introduce the method using the Cormack-Jolly-Seber model (CJS; Cormack, 1964; Jolly, 1965; Seber, 1965) for inference about the survival and abundance of marked animals under conditions of imperfect detection. The contribution of this paper is to make two modifications to the conventional component-wise boosting algorithm (e.g. Schmid et al., 2010) in order to make boosting appropriate for serially-dependent time-series of CMR data, a.k.a. capture-histories. One CJSboost method interweaves an Expectation-Maximization (EM) step between boosting iterations, and the second method uses stochastic imputation of latent states. Both methods can be used to estimate the *gradient of the loss function* which is the crux of statistical boosting. This paper is meant to prove and motivate these modifications so that boosting can be introduced to a wider-class of CMR models, such as the POPAN or PCRD or spatial capture-recapture. Code is available on the Github site http://github.com/faraway1nspace/HMMboost as well as a tutorial.

In this article, I introduce CJSboost by positioning it within the general theory of model-selection and multi-model inference (MMI); specifically, I show that CJSboost can be used for the two domains of multi-model inference: i) efficient estimation and/or prediction, and ii) consistent model-identification a.k.a. finding the hypothesis-cum-model which most support. These are what Aho et al. (2014) refers to as A-type vs. B-type thinking. I show why boosting is very appealing, both theoretically and practically, for CMR practitioners who use MMI techniques, such as AIC model-averaging or BIC model-selection.

Specifically, CJSboost is a type of *shrinkage estimator*: it negotiates the complexity of a model in order to minimize a prediction error. This error is closely related to the Expected Log-Likelihood which Akaike used to motivate his famous derivation of the AIC (Akaike, 1974, 1998). Akaike explained that model-selection according to the Expected Log-Likelihood is efficient: it performs best at minimizing the square-error between estimates and a true process. Through simulation, I show that boosting is qualitatively similar to AICc-methods at estimating abundance, and it is much better at estimating survival. I also propose that CJSboost can be coupled with a new technique called stability selection (Meinshausen & Bühlmann, 2010) in order to derive a *sparse estimator*, that is, to find covariates that significantly influence survival and are part of the “true model”, much like the BIC. Therefore, CMR practitioners can use the two flavours of CJSboost in order to tackle both domains of MMI: efficient estimation or consistent model-identification.

However, CJSboost has many other advantages over AIC/BIC model-selection and their constituent fixed-effect models:

- it can automatically perform variable-selection and explore higher-order interactions, even in situations of low-sample size (i.e., the *n < p* problem);
- it can include non-linear effects such as splines, regression trees, spatial kernels, or any of the base-learners available in the mboost family of R packages (Bühlmann & Hothorn, 2007; Hothorn et al., 2006; Mayr et al., 2012; Hofner et al., 2012);
- it has shrinkage of estimates away from extreme values and inadmissible values (e.g., 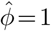) and avoids parameter singularities;
- its shrinkage properties can handle parameter non-identifiability issues better than the use of arbitrary constraints (e.g., fixing *ϕ*_*T*_ = *ϕ*_*T*–1_);
- it can better cope with multi-collinearity;

There are, however, many disadvantages and challenges to CJSboost. Some challenges are technical and require further research, such as theoretical validation of the consistency of stability selection. Other challenges are conceptual and will require practitioners to embrace new ideas and re-think old habits (such as reliance on p-values). I will briefly comment on some of the conceptual challenges first, then I will suggest new lines of research to address some technical challenges and useful extensions.

### 4.1. Conceptual challenges

Component-wise boosting is related to many important statistical ideas (Meir & Rätsch, 2003). It is similar to the Lasso solution (Efron et al., 2004; Bühlmann & Hothorn, 2007), which is favoured in machine learning. It is a type of model-averaging (Hand & Vinciotti, 2003) by weighting the outputs of hundreds or thousands of sub-models. It is also a Generalized Additive Model which is itself a type of penalized regression approach (Mayr et al., 2012). Despite these connections with other popular techniques, the ecological community has been slow to adopt statistical boosting. I believe this may be due to a few conceptual misunderstandings, such as shrinkage and suspicion of algorithmic learning techniques.

#### Algorithmic Inference

Boosting originally arose as a purely algorithmic means of classification (Meir & Rätsch, 2003; Mayr et al., 2014). Some ecologists have embraced such methods (Elith et al., 2008), but I suspect many are sceptical of machine-learning methods in favour of parametric Maximum Likelihood Estimation (MLE), especially given the long-studied optimality properties of the latter. Part of the motivation of this article was to review some theory about model-selection, such as shrinkage and Akaike’s AIC, and show why they lend support to component-wise boosting for statistical inference. Namely, we now know that at finite sample sizes, the MLE solution of a multiple-regression problem is inadmissible because of shrinkage (*sensu* Copas, 1983, 1997). Secondly, Akaike (1974) showed us that the Expected Log-likelihood, rather than the Maximum Likelihood, is efficient at deciding the optimal complexity of a model. Therefore, there is solid theory to support the statistical utility of CJSboosting for CMR analysis, given that it is a type of shrinkage estimator and it approximates the Expected (negative) log-Likelihood.

#### Shrinkage

Despite a huge body of research about shrinkage (Stein, 1956; James & Stein, 1961; Copas, 1983, 1997; Royle & Link, 2002), shrinkage creates a conceptual discomfort for ecologists, and this may be boosting’s greatest hurdle. First, we must do away with familiar tools like p-values and confidence intervals (more below). More importantly, we must grapple with the red-herring of unbiased-ness, to which most practical ecologists seem to consider sacrosanct. Ecologists trained to scrutinize diagnostic residual-plots may look at the bias in Figure 2 and be very alarmed, despite the underlying loss-optimality. In other words, we incur some bias to minimize an expected square-error loss (see Appendix E). This made shrinkage highly controversial 50 years ago at the time of its discovery (Efron & Morris, 1975), and its repercussions have not fully permeated the non-statistician research community.

#### Bayesian Interpretation

However, the rising popularity of Bayesianism may be the greatest advocate for component-wise boosting. First, *𝓁*_1_-regularizers, such as the Lasso and component-wise boosting, have a Bayesian interpretation (Geman et al., 1992; Hooten & Hobbs, 2015), and the outputs are merely a type of a Maximum A Posteriori (MAP) estimate (Murphy, 2012c). Secondly, ecological practitioners seem unconcerned with the fact that Bayesians are technically biased due to the role of priors at finite sample sizes. To wit, Bayesians have become popular champions of shrinkage, to the extent that it almost seems like a Bayesian idea, despite its Frequentist origins. For example, Royle & Link (2002) advocated for Hierarchical Bayesian random-effect models for CMR primarily because of the benefits of shrinkage. CJSboost is the Frequentist answer to their work.

### 4.2. Inference without Confidence Intervals or P-values

In this paper, I have chosen not to show 95%CI nor classical p-values for marginal effects’ null-hypothesis tests. I ignore these in order to focus the reader’s attention on point-wise estimation: the type of inference that shrinkage and AIC-like estimators were specifically developed for and should do optimally. For example, if one desires a time-series of abundance, then boosting or AIC-methods should produce estimates that generally have the lowest mean square-error loss between truth and estimate, i.e. the point-estimates are as close as possible to the truth, over all possible samples from the population. This type of inference does not depend on significant effect sizes or 95%CI; estimation variance is directly incorporated into the procedure through shrinkage (Appendix E). That being said, it is common in the boosting literature to use bootstrapping to approximate CI, and this could be done in CJSboost by bootstrapping capture-histories.

However, I would urge practitioners to think carefully about why they wish to have p-values or CI, rather than consider them as default statistics. There is growing concern about the misuse of p-values (Anderson et al., 2000; Gerrodette, 2011) and CI (Hoekstra et al., 2014), and some journals have started banning them altogether (Trafimow & Marks, 2015). I suggest that there are alternative tools which are more aligned with one’s research goals. For example, if a practitioner is interested in using 95%CI or classic p-values to test whether a covariate is “significantly” different from zero, then perhaps the real intention is to discover which covariates are truely influential? For this type of model-identification inference (what Aho et al., 2014, called B-type thinking), I propose the use of stability selection and approximate posterior inclusion probabilities. Similarly, one may wish to cap their False Discoveries (Meinshausen & Bühlmann, 2010; Shah & Samworth, 2013). This is a closer marriage of research goals and statistical analysis.

Finally, I would also remind readers that the abandonment of CIs or p-values is not a unique deficiency to CJSboost, but is true for all model-selection or shrinkage estimators. The common practice of doing model-selection and then using the CIs or classic p-values from the best model, as if model-selection was never performed, is invalid. Breiman (1992) called this a “Quiet Scandal”. The sampling properties of a post-model-selection estimator can be significantly different from those of a single-model (Leeb & Pötscher, 2005). This is the price of multi-model inference vs. declaring a true model *a priori*. Therefore, one’s only recourse in MMI is to use model-averaged CIs (Anderson et al., 2000) or bootstrap-approximated CIs, or multi-model p-values (Lee & Boone, 2011) or, better yet, to calculate statistics which actually address one’s research question.

### 4.3. Extensions and Future Considerations

This study is merely the first step in developing and introducing boosting for CMR models. A lot of the theory of loss-efficiency and consistency in univariate boosting for will need further validation in the HMM context.

#### Estimation

Regarding estimation performance, the simulations showed that CJSboost is very competitive, and perhaps better, than AICc averaging or model-selection at estimating survival and abundance. However, it is unknown whether CJSboost shares any of the theoretical efficiency properties of its univariate version. For example: does it obtain the minimal worst-case error, i.e. is it minimax optimal (Bühlmann & Yu, 2003)? How sensitive is its performance to the regularization parameters? Of more practical concern, the new basis functions of mboost create new ways to address old CMR estimation challenges, such as random-effect base-learners to accommodate individual heterogeneity, or CART for automatic discovery of non-linear processes. These opportunities require further empirical study, such as whether they incur significant estimation trade-offs. For example, Bühlmann & Yu (2003) found worse estimation performance with CART-like learners vs. least-square learners in simple linear regression models.

#### Consistency

Regarding variable selection or hypothesis-testing, this type of inference has been much less important in CMR than estimating abundance. However, I expect that it will become more important in certain “Big Data” domains where interest lies in finding significant associations between demographic variation and environmental covariates. For such inferences, the key property that a researcher needs is model-selection consistency: she desires a procedure that can recover the true model with high-probability. This type of MMI is prone to False Discoveries, especially when practitioners use prediction-optimal methods, such as the AIC/c or its derivatives (Shao, 1993; Yang, 2005). This misuse is widespread in ecology, and may contribute to the current crisis of reproducibility (Galipaud et al., 2014). For consistent variable selection, boosting has many potential extensions, such as TwinBoosting (Bühlmann & Hothorn, 2010). I suggest enhancing CJSboost with stability selection to approximate Bayesian inclusion probabilities.

#### Stability Selection

This is an exciting and growing field of study, and the stability-selection-enhanced CJSboost technique may need revision in the near future. In particular, the univariate versions of stability selection have theoretical bounds on the number of False Discoveries (Meinshausen & Bühlmann, 2010; Shah & Samworth, 2013) and selection probabilities of spurious variables (Bach, 2008), but these do not apply to multi-parameter boosting. Secondly, it is unclear whether we must marginalize over all three regularization parameters (*m* and *ν*_*p*_ and *ν*_*ϕ*_) or whether we can, as I have suggested, focus only on *m*. Third, it is unclear whether there is a violation of the assumption “exchangeability” of spurious covariates, as may be the case with the time-varying covariates vs. individually-varying covariates, as suggested in the simulations. These will require more empirical study. The latter may be partially solved by using the less-restrictive complementary-pairs stability selection of Shah & Samworth (2013). Nonetheless, the simulation results are promising and in-line with other studies: that is, influential covariates are selected with a probability that converges to 1 as sample sizes get large, and there is good discrimination between significant and negligible covariates.

#### Extensions

By validating the boosting technique for a simple open-population model, this study paves the way for more popular CMR models, such as POPAN and the PCRD, which have more model parameters in the likelihood function, like temporary-migration processes. With more parameters, the boosting algorithms will require more efficient ways of tuning regularization parameters. See Appendix C.2 for ideas in this regard.

#### New Base-learners

One major benefit of the CJSboost framework is its extensibility. It can accommodate phenomena such as individual heterogeneity, spatial capture-recapture and cyclic-splines. These are possible because the CJSboost code is written for compatibility with the `mboost` family of R packages, and leverages their impressive variety of base-learners (Bühlmann & Hothorn, 2007; Hofner et al., 2012). For example, the `brandom` base-learner can accommodate individual random effects for addressing individual heterogeneity in a manner similar to Bayesian Hierarchical models (Rankin et al., 2016). Kernels (`brad`) and spatial splines (`bspatial`) can be used for smooth spatial effects (Kneib et al., 2009; Hothorn et al., 2010; Tyne et al., 2015) offering an entirely new framework for spatial capture-recapture. The largest advantage is that users can add these extensions via the R formula interface, rather than having to modify deep-level code.

## 5. Conclusions

1. Boosting is a shrinkage estimator and regularization algorithm that can be adapted to capture-mark-recapture through an additional Expectation-Maximization step that imputes latent states.
2. Boosting negotiates the “bias-variance trade-off” by incurring a slight bias in all coefficients, but yields estimates that are more stable to outliers and over-fitting, across multiple realizations of the data (Appendix E).
3. CJSboost allows for powerful learners, such as recursive-partitioning trees (e.g., CART) for automatic variable-selection, interaction detection, and non-linearity. This flexibility seems to come at the cost of slightly more conservative estimates (if the underlying true model is linear).
4. Both AICc model-selection and boosting are motivated by good predictive performance: minimizing an expected loss (a.k.a. risk, or generalization error). When using least-squares or CART-like base-learners, the estimates from CJSboost are qualitatively similar to AICc model-averaging, but with more shrinkage on coefficients.
5. CJSboost seems to perform very well in high-dimensional model-selection problems, with the ability to recover a small set of influential covariates.
6. If the goal of a CMR analysis is to not estimate abundance or survival, but to find significant covariates, then CJSboosted models can be enhanced with stability-selection to derive a model-selection consistent estimator. Further research is necessary to validate the consistency property.

## 6. Acknowledgements

I would like to thank Professor Sayan Mukherjee for giving this project an initial “thumbs up” during a Duke University course on Probabilistic Machine Learning. I would also like to thank David Anderson, Lars Bejder, Krista Nicholson and Julian Tyne for helpful comments and critiques which greatly strengthened this manuscript.

## APPENDICES

### Appendix A. The CJSboost algorithm for Monte-Carlo approximation

The second strategy to boost a CJS capture-recapture model is called CJSboost Monte Carlo (MC). The idea is to garner conditional independence of data-points (*y*_*j*_, **x**_*j*_) by integrating over the distributions of latent states *π*(**z**_*i*_*|***y**_*i*_, **F**_*i*_). The integration is approximated with a large sample from the posterior of **z**_*i*_. A fast and simple “forward-filtering and backward-sampling” algorithm is used to sample latent states (Rabiner, 1989; Murphy, 2012b), detailed in Appendix B.4.

Within each boosting iteration *m*, we sample *S* sequences of **z**_*i*_. Per *s* sequence, we estimate a separate negative-gradient, and fit base-learners to it. After fitting all *S* samples, we update the prediction vectors with the best-fitting base-learners from each sequence, 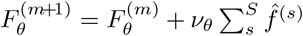. Over *S × m* draws, this is approximately equivalent to the EM algorithm. For comparable results to CJSboost-EM, the learning-rate parameters *ν*_MC_ should be set equal to 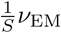, i.e., the contribution of any one sequence **z**^(*s*)^ is small.

I now describe the CJSboost-MC algorithm:

1. Set regularization parameters *S*, *m*_stop_, *ν*_*ϕ*_, and *ν*_*p*_.
2. Initialize *m* = 1 and 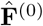.
3. For *s* = 1: *S*, do:

a. sample latent state sequence 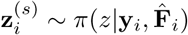 (see Appendix B.4);
b. estimate the negative gradients, conditional on 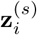:

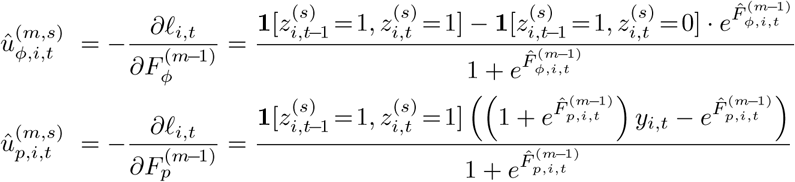
c. for each *θ* in {*ϕ, p*} do:

i. for each *k* base-learner in *θ* do:

A. fit the base-learner to the gradient: 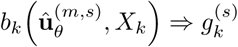;
B. make an estimate of the gradient, 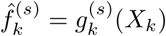;
ii. find the base-learner that best-fits the gradient 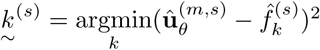;
iii. append the prediction function of 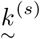 to the ensemble 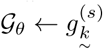;
4. Update the fit vectors for each *θ ∈* {*ϕ, p*}, taking the sum over all *S*: 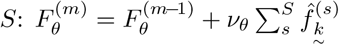.
5. Estimate the empirical risk 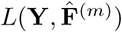, or estimate the holdout-risk on an out-of-sample subset of the data 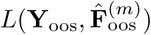 for cross-validation.
6. *m* = *m* + 1
7. Repeat steps 3 to 6 until *m* = *m*_*stop*_.

Just as in the CJSboost-EM algorithm, we must tune *ν* and *m*_stop_ through cross-validation or bootstrap-validation (Section 2.2.3).

Notice that the two algorithms have different surrogate loss functions and negative-gradients. However, the expected loss is still the Expected negative CJS Log-Likelihood, and the empirical risk is the negative CJS log-likelihood of the observed data.

Figures A.9 and A.10 compare the CJSboost-MC algorithm against the CJSboost-EM algorithm. Figure A.9 shows model estimates of capture-probability and survival for an example dataset from Simulation 1 of the main article; we see that the MC algorithm produces approximately similar estimates, although there is some extra variation in the *b*_trees_ base-learners model. Figure A.10 is from the high-dimensional Simulation 3 in the main article. The Figure shows a scatter-plot of the estimates from the EM algorithm vs. the MC algorithm, using a simulated high-dimensional dataset, where each dot is an individual *i* at capture-period *t*. The results fall along the 1:1 line, which demonstrates that the algorithms are approximately equivalent.

**Figure A.9:**
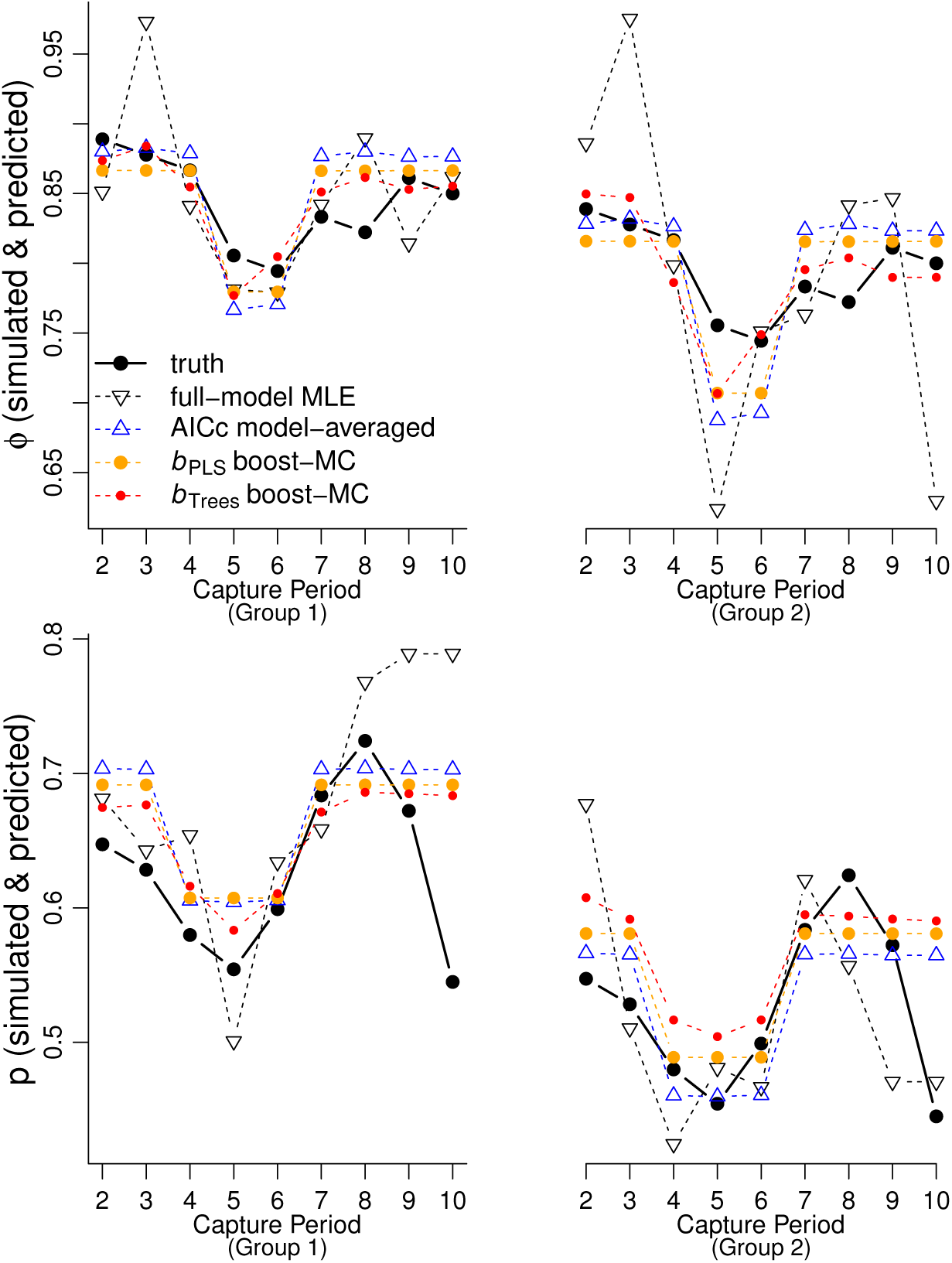
From Simulation 1 of the main article, a demonstration of CJSboost estimates from the Monte-Carlo approximation technique. A comparison of capture-probability estimates 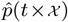 and survival estimates 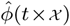 from four models: CJSboost-MC with linear base-learners (OLS and PLS; in **orange**); CJSBoost-MC with non-linear base-learners (CART-like trees; in **red**); AICc model-averaging (**blue**); and MLEs of the full-model (dashed black).

**Figure A.10:**
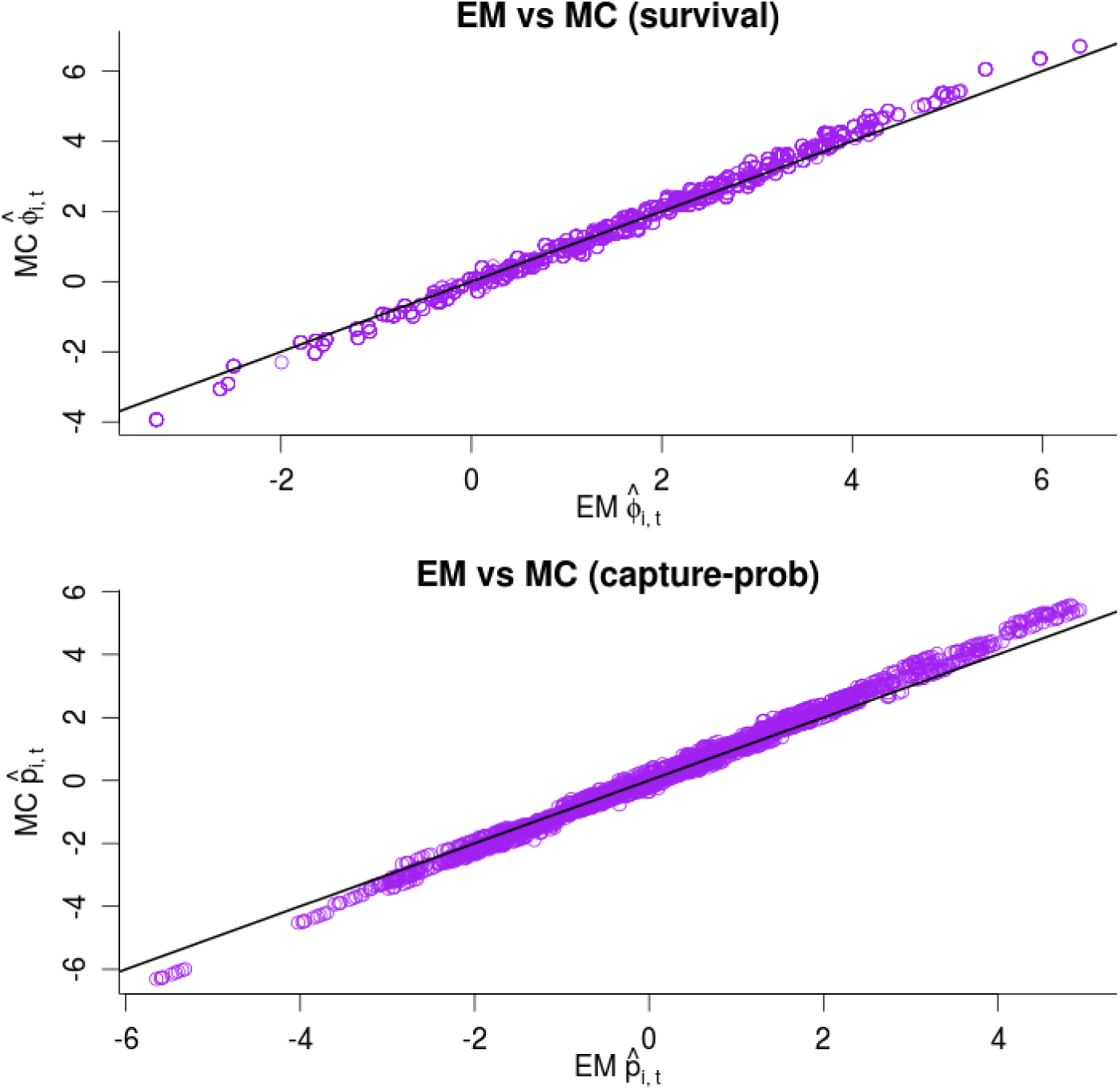
Simulation 1, demonstrating CJSboost estimates from the Monte-Carlo approximation technique. A comparison of capture-probability estimates 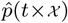 and survival estimates 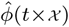 from models composed of linear base-learners (OLS and PLS; in orange) and non-linear base-learners (CART-like trees; in red), as well AICc model-averaging (blue) and MLE (dashed black).

### Appendix B. Algorithms for Filtering and Sampling HMM Latent States

The CJSboost algorithms depend on conditional independence of data pairs (*y*_*i,t*_*, X*_*i,t*_) for individuals *i* in capture period *t*, in order to estimate the negative-gradient in the descent algorithm. This is possible if we impute information about the latent state sequences *z* for pairs of capture periods at *t* and *t* – 1. The two CJSboost algorithms, CJSboost-EM and CJSboost-MC, achieve this same idea with two different, but related, techniques. In both cases, we will use a classic “forwards-backwards” messaging algorithm to gain information about the probability distribution of the latent state sequences. In CJSboost-EM, we calculate the *two-slice marginal probabilities p*(*z*_*t*–1_ = *u, z*_*t*_ = *v|***y**_1:*T*_ *, ϕ, p*), per boosting iteration; in CJSboost-MC, we will *sample* **z** from its posterior distribution *π*(**z**_1:*T*_ *|***y**_1:*T*_ *, ϕ, p*). See Rabiner (1989) and Murphy (2012b) for accessible tutorials.

Both algorithms use a forwards-messaging algorithm and a backwards-messaging algorithm. The forwards algorithm passes information about the state of *z*_*t*_ conditional on all previous observations (denoted *α*_*t*_), whereas the backwards algorithm estimates the future conditional likelihood of the capture-data given *z*_*t*_ at *t* (denoted *β*_*t*_). The *α* and *β* values are combined to make inferences about the distribution of latent states per time *t*.

We will drop the indices *i*, and focus on the capture-history of a single individual. **y** is the time-series of binary outcomes of length *T*. **z** is a vector of latent states *z ∈* {dead, alive}. We condition on an individual’s first capture at time *t* = *t*^*0*^, and are only concerned with the sequence **z**_*t*^0^:*T*_. Survival from step *t*–1 to *t* is *ϕ*_*t*_. Conditional on *z*_*t*_, the capture-probabilities are *p*(*y*_*t*_ = 1*|*alive) = *p*_*t*_, and *p*(*y*_*t*_ = 1*|*dead) = 0. In HMM notation, the CJS processes can be presented as the following column-stochastic matrices:

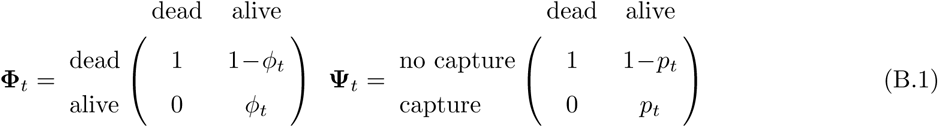

In HMM parlance, **Φ** is the Markovian transition process; we denote the probability *p*(*z*_*t*_ = *u|z*_*t*–1_ = *u*) as **Φ**_*t*_(*u, v*). **Ψ** is the emission process representing the conditional capture-probabilities; we denote the probability *p*(*y*_*t*_ = 1*|z*_*t*_ = *v*) as **Ψ**_*t*_(*v*).

#### Appendix B.1. Forwards-algorithm

The forward messaging algorithm involves the recursive calculation of *α*_*t*_(*v*), per time *t* and state *z*_*t*_ = *v*. *α*_*t*_ is the *filtered belief state* of *z*_*t*_ given all the observed information in **y** from first capture *t*^0^ until *t*. Notice, that for clarity, we drop the notation for conditioning on *ϕ* and *p*, but these are always implied.

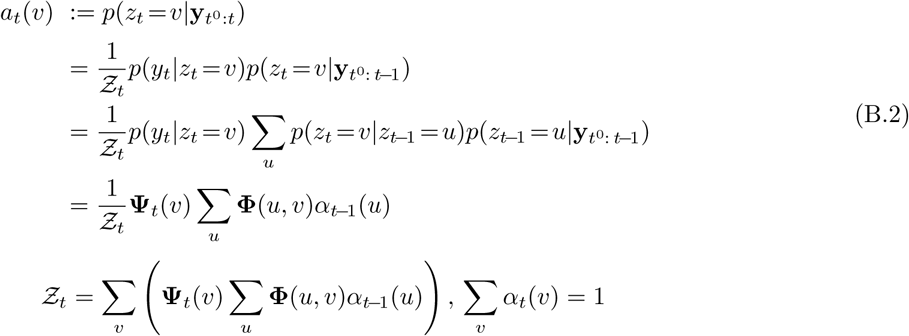

The algorithm is initialized at time *t*^0^ (an individual’s first capture) with *α*_*t*_0 (alive) = 1 and *α*_*t*_0 (dead) = 0. This is true because the animal must be alive for us to capture it. Conditional on the values of *α*_*t*_(*v*) for all *v*, one can proceed to calculate the next values of *α*_*t*+1_(*v*), and so on, until *t* = *T*.

#### Appendix B.2. Backwards-algorithm

Messages are passed backwards in a recursive algorithm starting at *t* = *T* and moving backwards until *t* = *t*^0^, the first-capture period, while updating entries in *β*_*t*_(*v*). *β*_*t*–1_(*u*) is defined as the likelihood of future observations **y**_*t*: *T*_ from *t* to *T*, conditional on *z*_*t*–1_ = *u* at *t*–1.

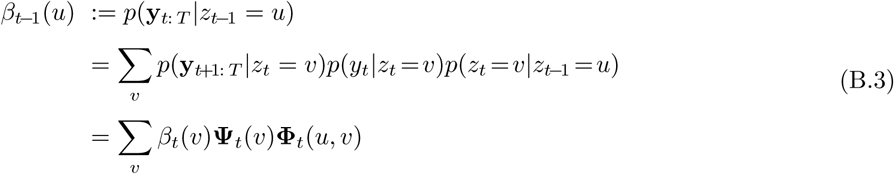

The algorithm is initialized *β*_*T*_ (·) = 1 for all states *v*, and proceeds backwards as above. Notice that the elements of *β*_*t*_(·) do not need to sum to 1.

Having calculated the backwards and forwards messages, we can now proceed to characterize the latent state distributions and boost *ϕ* and *p*.

#### Appendix B.3. Two-slice marginal probabilities for Expectation-Maximization

Expectation-Maximization is an iterative technique for maximizing a difficult objective function by working with an easy “complete-data” objective function log *p*(*y, z|θ*). EM works by cycling through an M-step and an E-step. In boosting-EM, the M-step corresponds to the usual update of the fit vectors 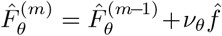 (conditional on *z*), which are used to estimate 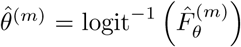. The E-step corresponds to imputing the expectations of the latent states *z*, conditional on the data and current estimates of 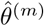.

Technically, we require the expectations for the *pairs* of sequential states (*z*_*t*–1_*, z*_*t*_). In CJS, these pairs of states are simply {*alive, alive*}, {*alive, dead*}, {*dead, dead*}. Using the Complete-Data Likelihood, we substitute in the two-slice marginal probabilities *w*_*t*_:= *p*(*z*_*t*–1_*, z*_*t*_*|***y**_*t*^0^_:*T, ϕ, p*) for the pairs (*z*_*t*–1_*, z*_*t*_). These probabilities can be calculated easily for a capture-history **y**_*i*_ using the outputs (*α, β*) from the forward-backwards algorithm.

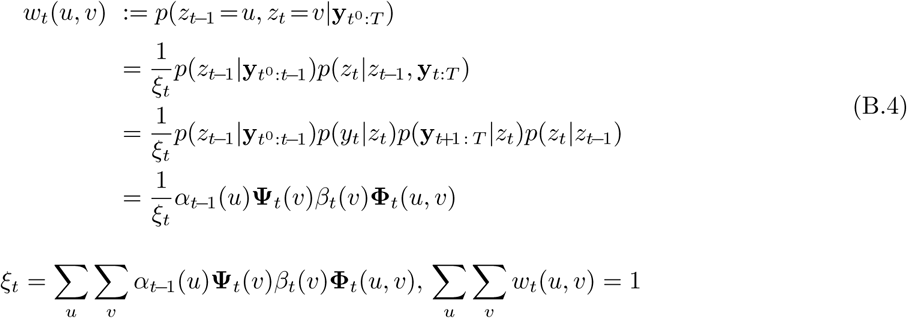

The E-step is completed after evaluating the set {*w*_*i,t*_(alive, alive)*, w*_*i,t*_(alive, dead)*, w*_*i,t*_(dead, dead)}, for each capture period 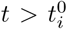 and for each individual 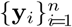. This is an expensive operation; computational time can be saved by re-evaluating the expectations every second or third boosting iteration *m*, which, for large *m*_stop_ *>* 100 and small *ν*, will have a negligible approximation error.

#### Appendix B.4. Sampling state-sequences from their posterior

For the CJSboost Monte-Carlo algorithm, we sample a latent state sequence **z**_*i*_ from the posterior *π*(**z**_1:*T*_ *|***y**_1:*T*_ *, ϕ, p*), for each individual *i* per boosting step *m*. Conditional on the latent states, the negative-gradients are easily evaluated and we can proceed to boost the estimates and descend the risk gradient. However, because the algorithm is stochastic, we must avoid getting trapped in a local minima by sampling many sequences (e.g., *S ≈* 10 – 20), thereby approximating the full posterior distribution of **z**. Over all *S* samples, the average gradient will *probably* be in the direction of the global minima. For large *m* and small *ν*, the approximation error is small.

The algorithm performs backwards-sampling of the posterior using the chain rule:

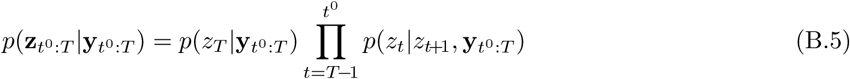

We start with a draw at time 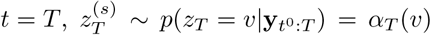, and condition earlier states on knowing the next-step-ahead state, proceeding backwards until *t* = *t*^0^.

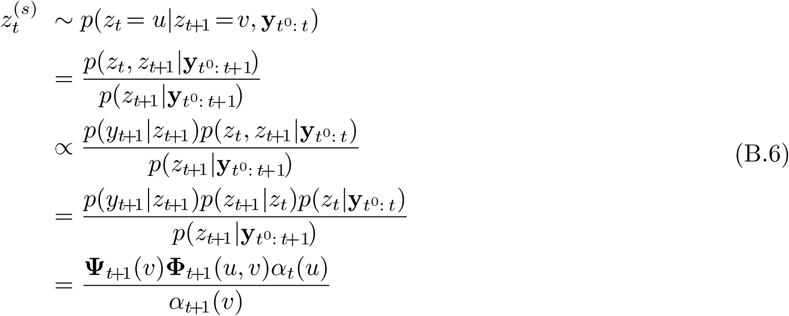

Thus, knowing *α*, *β*, **Φ** and **Ψ**, we can easily generate random samples of **z** from its posterior distribution. The backwards sampling step is repeated for each 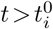 capture period, for each *s* sequence, for each individual *i*, and for each *m* boosting iteration.

### Appendix C. Algorithms for Tuning the Regularization Parameters

This section will present a simple work-flow for finding approximately optimal values of *m*_*stop*_, *ν*_*ϕ*_ and *ν*_*p*_ that minimize our expected loss *ℒ*, a.k.a. the generalization error. We approximate *ℒ* through *B*-fold bootstrap-validation. For each *b* bootstrap, we create a CJSboost model, *G*^(*b*)^(*X*; *m, ν*_*ϕ*_*, ν*_*p*_) which is trained on the bootstrapped data and is a function of the regularization parameters *ν*_*ϕ*_, *ν*_*p*_ and *m*. We calculate the holdout-out risk using the out-of-bootstrap *b*^*c*^ capture-histories and covariate data, (**Y**^(*b*^*c*^)^, **X**^(*b*^*c*^)^). The objective to minimize is the average hold-out risk, *L*_cv_, estimated over *B* bootstraps.

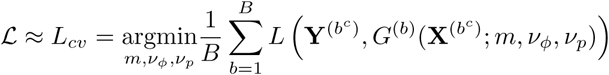

In univariate boosting, it is easy and routine to find the optimal *m*_stop_ through bootstrap-validation, conditional on a fixed value of *ν*. It is easy because we can simultaneously fit a model *and* monitor the holdout-risk per *m* step. Therefore, we need only perform one round of bootstrapping to find the *m*_cv_ that minimizes the average holdout-risk.

However, the focus of this section will be to estimate the optimal values of *ν*_*ϕ*_ and *ν*_*p*_. This is a seemingly difficult task because they are continuous: we cannot realistically run a different bootstrap exercise per combination of ℝ^+^*×* ℝ^+^. The challenge of optimizing *ν*_*p*_ and *ν*_*ϕ*_ is not unique to CJSboost, but is inherent to all multi-parameter boosting techniques, such as boosted-GAMLSS. Readers who are already familiar with the boosted-GAMLSS literature may notice that my approach differs slightly from other authors (e.g. Schmid et al., 2013; Mayr et al., 2012). These authors used a single fixed value of *ν* for all parameters, and then optimized separate values of *m*_*θ*_ per parameter *θ*. Alternatively, I propose to optimize a global *m*_stop_ for both parameters, after optimizing the *ratio* of *v*_*θ*_1__ to *v*_*θ*_2__. The two methods are equivalent in their outcome. I wish to emphasize that although the boosting literature has claimed that there is little benefit in optimizing *m* and/or *ν* separately for each parameter (Schmid et al., 2013), this is untrue for CJSboost. The optimal estimate of 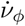 may be several orders of magnitude different than the optimal 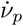.

The most easy-to-understand method to optimize *ν*_*ϕ*_ and *ν*_*p*_ is to discretize the set of plausible combinations, such as (10^–4^, 10^–3^, 10^–2^, 10^–1^) ⊗ (10^–4^, 10^–3^, 10^–2^, 10^–1^). This is not a terrible idea because Bühlmann & Yu (2003) showed that the generalization error has a very shallow minima around the optimal values of *m*. This means that our regularization parameters need only get within the vicinity of their optimal values, rather than strict numerical convergence. However, searching for optimal values on a small grid of combinations would be very expensive and imprecise. Therefore, we seek an adaptive algorithm that can get closer to the optimal values of *ν*_*ϕ*_ and *ν*_*p*_ with only 7-10 bootstrap-validation exercises.

#### Appendix C.1. Algorithm 1 for Optimizing Learning-Rates

For just two parameters (*ϕ*,*p*), we can find the minimum *L*_cv_ by optimizing the ratio 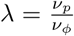, for a fixed mean 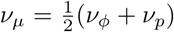. We can safely fix *ν*_*μ*_ because it has a straight-forward inverse relationship to *m*_stop_; so if we fix one, we merely solve for the other. The point is that we have reduced the problem to a univariate search to find the 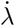 which minimizes *L*_cv_(*λ*). Recall also that we can always find the optimal *m*_stop_ for a given *λ* and *ν*_*μ*_, so we can drop *m* from our objective function, which is now a univariate objective:

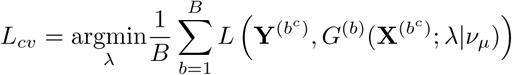

This is less daunting than it may seem, because the range of *λ* is practically bounded. For example, for large *m*_stop_ and 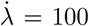, then *ν*_*p*_ *≫ ν*_*ϕ*_, and *ϕ* is effectively shrunk to its intercept starting value. Higher values of *λ* will have little effect on the generalization error. Also, *L*_cv_(*λ*) is typically a convex function of *λ* (assuming that as we reuse the same bootstrap-weights for all new estimates of *L*_cv_(*λ*)). In other words, we are searching a U-shaped Real-line for its minimum. This means we can employ any convex optimization algorithm for a univariate non-differentiable function to iteratively search for the optimal 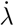.

The thrust of any such algorithm is a multiplicative “stepping-out” procedure to quickly find the correct order of magnitude for 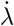. For example, starting at *λ*^(0)^ = 1, we need only 7 doubling steps to grow *λ* to 128 *× λ*^(0)^ ; further refinements will have little practical impact on the final model estimates. I suggest the following convex optimization algorithm:

1. set *ν*_*μ*_ = 0.01 and *λ*^(0)^ = 1; generate the *B* bootstrap samples and their out-of-sample compliments;
2. initialize the sorted list 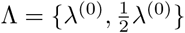;
3. for each *λ* in Λ, estimate *L*_*cv*_(*λ*) and store the values in the list 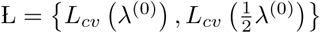;
4. for *j* in 1: *J*, do:

a. get the current best value for the ratio 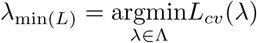
b. propose a new candidate *λ*\*: if *λ*_min(*L*)_ = min(Λ), then 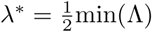 else if *λ*_min(*L*)_ = max(Λ), then λ* = 2.max(Λ) else *λ*\* = *λ*_min_ + *k* · *α*, where *k* is the step direction and *α* is the step size.
c. re-calculate the learning rates from 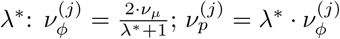
d. perform bootstrap-validation to estimate 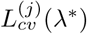;
e. append Λ *← λ*\* and append 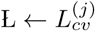;

The algorithm continues until a pre-defined convergence criteria is met, or, practically, a maximum number of *J* iterations is reached. The final values of *ν*_*ϕ*_, *ν*_*p*_, and *m*_cv_ are those which correspond to the minimum *L*_cv_ *∈* L.

There are many convex optimization algorithms which differ in how they calculate *k* and *α*. In CJSboost, most of the optimization benefits occur during the “stepping-out” procedure, and so exact values of *k* and *α* are less important, so long as they guarantee convergence. I suggest the following sub-algorithm (nested within step 4b above). This is entirely arbitrary but succeeds in quickly ruling-out large sections of suboptimal values of *λ*.

1. Define the triplet set Γ composed of the current best estimate of *λ*_min(*L*)_ as well as the sorted values just to the left and right, such that 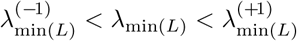;
2. Sort the entries of Γ according to the order *L*_*cv*_(*γ*^(1)^) *< L*_*cv*_(*γ*^(2)^) *< L*_*cv*_(*γ*^(3)^);
3. Estimate the step size and direction: if ∥*γ*^(1)^ *-γ*^(2)^ ∥ *≥* ∥*γ*^(1)^ *-γ*^(3)^ ∥: then 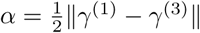 and *k* = sign(*γ*^(1)^ *-γ*^(2)^); else 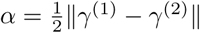 and *k* = sign(*γ*^(1)^ *-γ*^(3)^);
4. *λ*\* = *λ*_min(*L*)_ + *k* · *α*

Typically, seven or ten iterations are necessary in order to find suitable values of 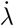, 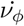 and 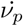. Unfortunately, this strategy is only useful for a two-parameter likelihood with a single ratio to optimize. For other capture-recapture models with more parameters (e.g., POPAN, PCRD), a different tuning strategy may be necessary, such as a bivariate convex optimization algorithm.

#### Appendix C.2. Algorithm 2 For Tuning the Learning-Rates ν

With more parameters in the capture-recapture likelihood, the number of necessary steps in algorithm 1 will increase exponentially. I suggest a second iterative algorithm whose number of iterations may only increase linearly with the number of parameters.

The principle of this second algorithm is based on the observation that when the ratio 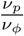 is poorly optimized, then additional boosting steps along the gradient 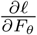 will *over-fit* and *increase* in the holdout-risk. This happen asymmetrically for *F*_*ϕ*_ vs *F*_*p*_. Therefore, we can monitor the extent of the asymmetry and adjust the ratio 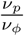 until the number of boosting steps which successfully decrease the hold-out risk is roughly the same for *F*_*ϕ*_ vs *F*_*p*_ (averaged over all bootstrap hold-out samples).

Call 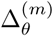 a boosting step along the partial derivative of 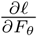 which successfully reduces the holdout-risk. I suggest using the ratio of Δ_*p*_ vs. Δ_*ϕ*_ as an estimate of 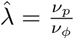.

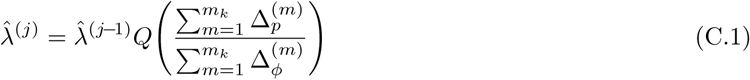

where *Q* is a robust measure of central tendency (e.g., trimmed mean) over all *B* bootstraps, and *m*_*k*_ is some boosting step *m*_*k*_ *≫ m*_cv_.

The first estimate 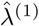 will typically be an underestimate, so the algorithm is iterated, each time using the previous values of 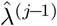 for setting 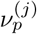 and 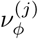 used to run CJSboost. The bootstrap-validation exercise is repeated to estimate the next 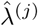 value according to Eqn. (C.1). 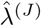will typically converge to a single value within approximately 10 iterations. 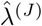is *not* the optimal 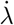 as estimated by algorithm 1, but it is within the vicinity of the optimal value (Figure C.11).

**Figure C.11:**
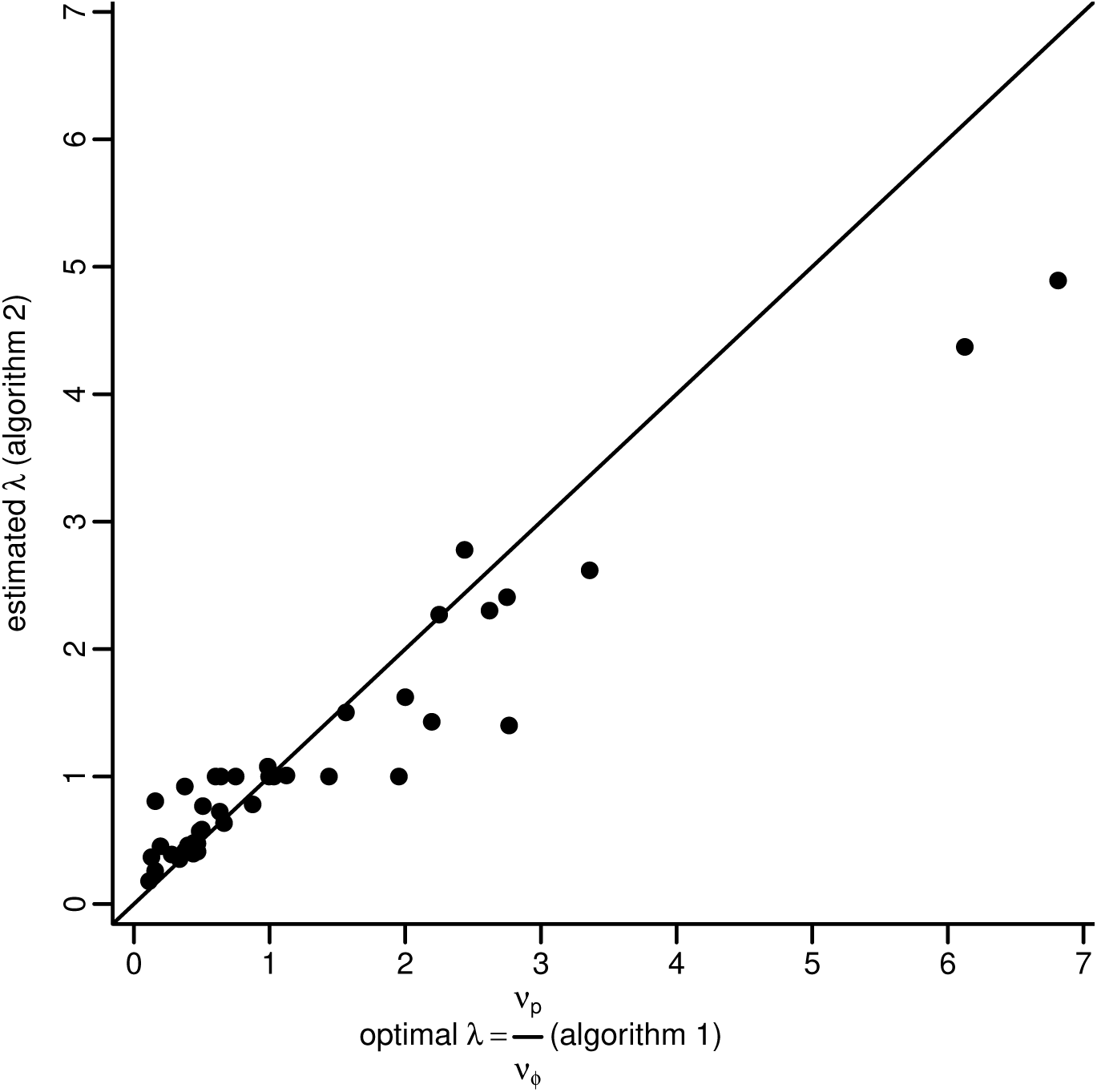
Two algorithms for tuning the learning-rate regularization parameters *ν*_*ϕ*_ and *ν*_*p*_, and their ratio *λ*, in order to minimize the expected loss (estimated via bootstrap-validation). Forty simulations compare the two algorithms, where algorithm 1 is considered optimal.

For just two *ν* parameters and one ratio (as in CJSboost), this second algorithm is not competitive with algorithm 1. But, when there are more than two parameters in the likelihood, this algorithm can simultaneously estimate all pertinent ratios.

Further refinements will be necessary. However, these preliminary simulations suggest that the risk gradient trajectories have information which can help optimize the regularization parameters.

### Appendix D. Specifying Base-learners

In component-wise boosting, there are some base-learner parameters that must be specified *a priori*. For example, PLS and P-spline base-learners have *effective degrees-of-freedom* parameters which constrain their flexibility to fit a process. Schmid & Hothorn (2008a) suggest that such parameters can be fixed to default values, and that practitioners should instead focus primarily on optimizing *m*_stop_. Furthermore, Bühlmann & Yu (2003) suggest that base-learners should be relatively weak, *a priori*, and that the overall model complexity should be tuned by controlling the shrinkage parameters *m*_stop_.

A more important consideration is the relative flexibility of competing base-learners. For example, multi-covariate learners and unpenalized learners have more flexibility to fit a process and minimize estimation error. Therefore, they may be preferentially selected in the component-wise boosting algorithm: recall that in step 7(b) of the CJSboost algorithm, it selects the best base-learner by a goodness-of-fit criterion. Therefore, practitioners should enforce a similar effective degrees-of-freedom among all base-learners, as well as decompose higher-order interactions and non-linear curves into their constituent components.

For example, if one desires model-selection among covariates *x*_1_ and *x*_2_ and their interaction *x*_1_ *× x*_2_, then one should specify four PLS base-learners of equal effective-*df*: one PLS base-learner for the *x*_1_ main-effect; a second PLS base-learner for the *x*_2_ main-effect; a third PLS base-learner for the main-effects of both *x*_1_ and *x*_2_ together (no interaction); and a final PLS base-learner for the interaction. This would be analogous to a shrinkage version of the R GLM model `glm(∼x1*x2,…)`. In the mboost R formula interface, the boosted model would be set-up with the following syntax:

~~~
∼ bols(x1,df=2)+bols(x2,df=2)+bols(x1,x2,df=2)+bols(x1,by=x2,df=2)
~~~

For non-linear splines on *x*_1_, we may wish to separate the linear and non-linear components, called “centring” in Kneib et al. (2009) and Hofner et al. (2012). In this case, the mboost formula interface would be `∼bols(x1)+bbs(x1,center=TRUE,df=1)`.

The above techniques are especially important if practitioners wish to gain some mechanistic understanding of the *ϕ* and *p* processes, such as concluding which covariates have a significant contribution to survival. This is crucial for using the stability-selection-enhanced CJSboost to find ecologically important covariates. However, when the research goal is not to uncover significant effects, but merely to accurately estimate abundance, then it is less important to enforce equal *effective-df* among base-learners. An extreme form of this is when estimation becomes a “black-box” exercise, for example, as with CART-like base-learners: `∼btree(x1,x2,tree_controls=ctree_control(maxdepth=2))`. Here, variable selection and non-linear effects and interactions are automatically incorporated, at the expense of interpretability.

### Appendix E. Primer On The Bias-Variance Trade-off

This appendix uses simulations to illustrate the “bias-variance trade-off” and shows how CJSboost and the AICc each negotiate the trade-off in order to minimize the expected error of estimating survival *ϕ* over T capture periods. The trade-off is fundamental to understanding the optimality of Frequentist shrinkage estimators and AIC model-selection. The illustrations are inspired by Murphy (2012a, figure 6.5), but adapted to capture-mark-recapture and the Cormack-Jolly-Seber model.

The trade-off is an old idea without a citable origin (although Geman et al., 1992, is often considered to be a definitive reference, but the phenomenon is clearly discussed as early as 1970 by Hoerl & Kennard). Despite being an old and fundamental concept of statistical estimation, I have noticed that it poorly understood among academics and government scientists. In particular, it is my experience that ecologists are unduly wedded to the idea of being unbiased (in estimation), such that when they are presented with visual and quantitative evidence about the optimality of biased shrinkage estimators, they recoil at the sight of systematic bias, and ignore the crucial role of variance. Of course, bias is not desirable in and of itself, but so long as the bias goes to zero at a rate proportional to that of the variance, we may be able to improve our overall estimation performance by incurring a little bias.

In the following simulations, the goal is to minimize the Expected Error of estimating survival, as quantified by the Mean Square Error (MSE). It is a population-level abstract quantity that can only be measured in simulations when we know to the “true” process. It is Frequentist in the sense that we hope to minimize the error over all possible data-sets that one might sample from the true population 𝕐. These multiple realizations are shown as grey lines in Figures E.12 and E.13. Of course, an analyst only has one dataset, and his goal is to get his estimates as close as possible to the truth.

**Figure E.12:**
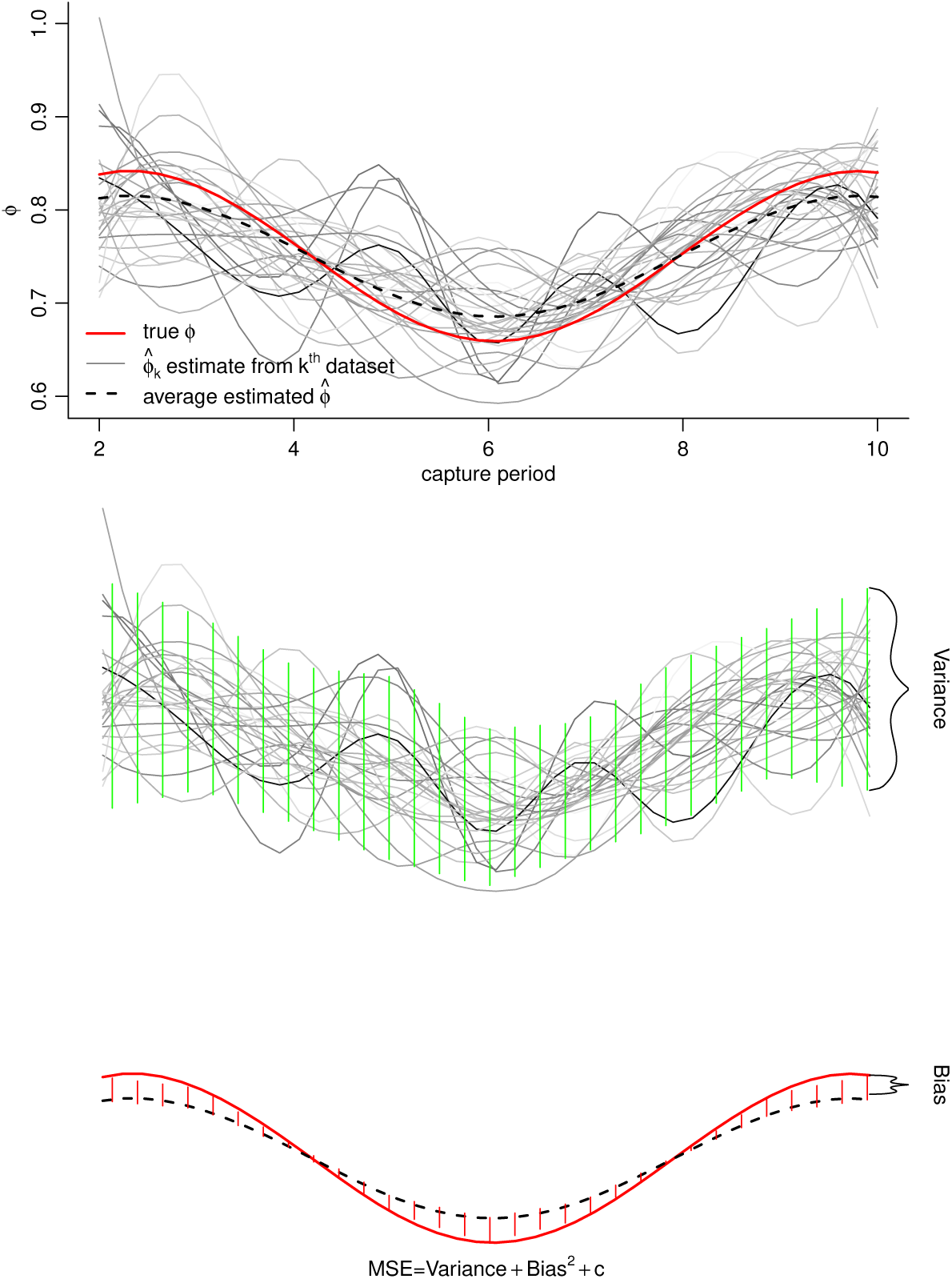
Decomposing the error of estimation (MSE) into its bias and variance components. An estimation procedure will negotiate the bias and variance so to minimize the MSE. *Top*, a simulation of a true survival process (red line). Each grey line represents one dataset sampled from the population and an analyst’s attempt to estimate survival using multi-model inference procedures, such as boosting. The dashed black line is the mean estimate over all 30 independent grey-lines. *Middle*, a visualization of the variance component, showing the variability of point-wise estimates due to randomness in the sampled data and a procedure’s sensitivity to such differences. *Bottom*, a visualization of the bias: the expected difference between the truth and the procedure’s estimates, over all realizations of the data.

**Figure E.13:**
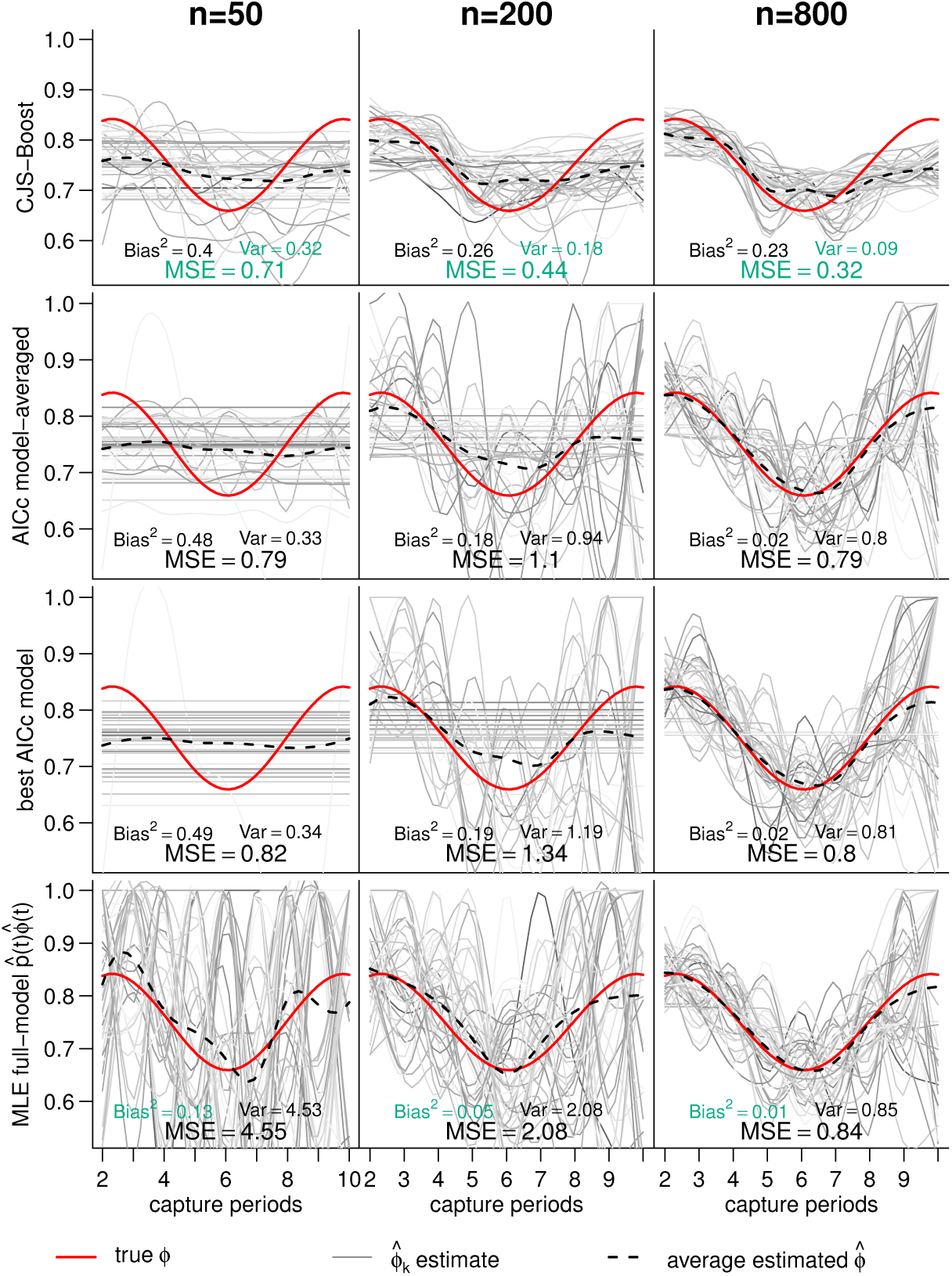
Visualizing the bias-variance trade-off and the error of estimating survival in a Cormack-Jolly-Seber analysis, using four procedures (*panel rows*): *i*) the shrinkage estimator CJSboost; *ii*) AICc model-averaging based on four fixed-effect models of time-varying vs. time-constant survival and capture-probabilities; *iii*) the best AICc model; and *iv*) the Maximum Likelihood Estimate using the full-model *p*(*t*)*ϕ*(*t*). *Panel columns* are different sample sizes (number of capture-histories) over *T* = 10 primary periods. The red-lines show the true survival. Each grey line is an independently sampled dataset and an analyst’s attempt to estimate survival. The dashed-lines represent each procedure’s average estimate over 40 simulated data-sets and analyses. The best estimation procedure has the lowest MSE (turquoise for emphasis). Each procedure may have a high/low bias or low/high variance, but generally cannot succeed at minimizing both. The bias is the difference between the red and dashed line. The variance is represented by the dispersion among grey lines. At small sample sizes, the AICc methods and boosting are very biased but have better MSE.

The bias-variance trade-off arises from a classic decomposition of the expected error: 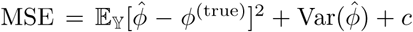. Figure E.12 also shows this decomposition. The first term is the expected difference between an estimate and the true value, i.e. the bias. This difference is visualized as the red polygon in Figure E.12. In the same figure, the bias manifests as shrinkage from the true red line towards a flat global mean. Quantifying the bias requires knowledge of the truth *ϕ*^(true)^, and is therefore inaccessible in real-life situations. The second term is the variance and it does not depend on knowledge of the truth. Rather, it arises due to the vagaries of random sampling as well as the complexity of the estimation procedure: overly complex models which “over-fit” one dataset will vary wildly when fitted to a new dataset sampled from the same population. The variance can be visualized as the spread of the grey lines, or the green polygon in Figure E.12.

The MSE decomposition has a naive meaning: in order to optimize our estimation performance, we should reduce the bias and/or the variance. Clearly, most ecologists see the value of tackling either of these two terms. But the nature of a *trade-off* has a more elusive importance: we cannot, in general, minimize both terms for a given sample-size, and we may deliberately increase one term in order to decrease the other. Shrinkage estimators incur a little bias and have lower variance (i.e., the red polygon is bigger but the green polygon is smaller). This strategy results in much smaller MSE values than complex unbiased estimators. In contrast, the MLEs of the complex full-model are unbiased but they typically have very high variance. This strategy is often worse at minimizing the MSE, for small-to-moderate samples sizes.

The following simulations show how different statistical methods have different strategies in negotiating the bias-variance trade-off. Imagine an analyst who is confronted with four different methods to estimate survival. The first is estimation by Maximum Likelihood using the full-model *p*(*t*)*ϕ*(*t*). The second method is AICc model-selection, and the third is AICc model-averaging; both use the following fixed-effects models: *p*(·)*ϕ*(·), *p*(*t*)*ϕ*(·), *p*(·)*ϕ*(*t*) and *p*(*t*)*ϕ*(*t*) with constraints on *p*_*T*_ = *p*_*T*–1_ and *ϕ*_*T*_ = *ϕ*_*T*–1_ terms. The fourth method is CJSboost with base-learners equivalent to the aforementioned fixed-effect models (but without the previous constraints). The AICc-methods should theoretically do best because they are fundamentally motivated by trying to minimize an objective function that is very closely related to MSE called the KL-loss (Akaike, 1974, 1998). Likewise, CJSboost is trying to minimize a related generalization-error called the negative Expected log-Likelihood, which is approximated through bootstrap-validation.

The fake data-sets were generated according to the following. 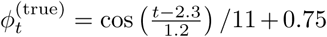. 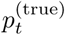 were drawn from a beta distribution with shape parameters *A* = 12 and *B* = 12, resulting in an average capture-probability of 0.5. The 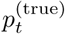 values were the same for all simulations. The first-captures were distributed randomly through-out the capture periods *t ∈* {1*, …,* 10}, with highest weight on *t* = 1. MLE and AICc analyses were run in Program MARK (White & Burnham, 1999) and RMark (Laake, 2013). For CJSboost, a ten-times 70-fold bootstrap-validation exercise was run per dataset to tune the CJSboost regularization parameters. The simulations and analyses were repeated 40 times for three scenarios pertaining to the number of capture-histories *n ∈* {50, 200, 800}.

The results clearly show the trade-off (Figure E.13). At high sample sizes (*n* = 800), the shrinkage estimator CJSboost has the lowest MSE and therefore wins at estimating survival. However, it has the highest bias. How can it be considered a better estimator than the other methods when it is biased? The answer is obvious when looking at the grey lines in Figure E.13, where each line is an estimate of 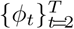 from an independent realization of data: compared to the other methods, each grey line from CJSboost is much more likely to be closer to the truth, despite systematic bias. In contrast, using the MLEs, one can only claim to be unbiased *over all possible realizations of the data* as shown by the closeness of the dashed black line to the true red line. But, for any one realization (a single grey line) the MLEs can be very far away from the truth due to much higher variance.

At smaller sample sizes, we see that the bias becomes much more extreme for both AICc methods and CJSboost. In the case of the AICc methods, the model with most support is often *ϕ*(·), in which case the estimates are a single flat line. This is also the case in CJSboost, were shrinkage is so extreme as to force a flat line. Therefore, at low sample sizes, we are much better off, in terms of MSE, to use the flat-lined 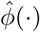 estimates rather than use the full-model MLEs, which vary so wildly as to be useless.

This primer is meant to illustrate the role of bias and variance in estimation errors. Simulations show how shrinkage estimators (CJSboost) and model-selection (by AICc) each negotiate the trade-off between bias and variance to try and minimize the Expected Error. CJSboost does particularly better by incurring a little bias.

### Appendix F. Extra Notes on Stability Selection

In the main article, I introduce stability selection for capture-mark-recapture (CMR) and use it to enhance the consistency properties of CJSboost, called SS-CJSboost. Stability selection is a new and rapidly growing group of methods, and SS-CJSboost borrows elements from different but related techniques by Bach (2008) and Meinshausen & Bühlmann (2010, hereafter referred to as *MeBü*) and Shah & Samworth (2013, *ShSa*). In this appendix, I will highlight how SS-CJSboost relates to these methods and where further validation may be necessary.

To review, the proximate aim of SS-CJSboost is to calculate 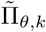, an approximation of the posterior inclusion probability, *π*(*β*_*θ,k*_ ≠ 0*|***Y**, **X**): the probability that a *k*^th^ covariate is part of the correct model of *θ*. Inclusion probabilities are routine in Bayesian analyses to address questions such as: does covariate *k* have some structural influence on survival? The analysis proceeds by bootstrapping the capture-histories *B* times, and for each *b* bootstrap running a CJSboost model on the *b*^th^ resampled data. We must score whether a covariate has been selected by CJSboost and has entered the ensemble 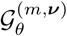 for each value of the regularization parameters (*m,* ***ν***) and for each *k* covariate and for each *b* bootstrap and for each *θ ∈* {*ϕ, p*}. We denote this selection indicator 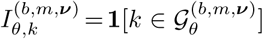. A short-cut is to pre-optimize the values of *ν*_*ϕ*_ and *ν*_*p*_, exactly as one would do in regular CJSboost analysis, and then condition all SS-CJSboost bootstrapped models on these values, called ***ν***?. The stability selection probabilities are calculated over *B* bootstraps per *m* and *k* and *θ*: 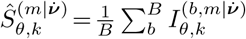. Finally, our Frequentist inclusion probability is the *mean* of the stability selection probabilities summed over all values of the regularization parameter 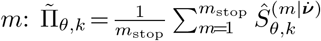.

Crudely, SS-CJSboost is most similar to the Bolasso (Bach, 2008), but with an emphasis on inclusion probabilities, as discussed in MeBü and ShSa. In the following paragraphs, I explain where and why certain techniques were incorporated into SS-CJSboost, and possible problems with the assumptions.

#### Selection Procedure

Bach, MeBü, and ShSa all demonstrate their methods on the Lasso. For Bach, the consistency results only hold for a region of the Lasso-regularization parameter in relation to sample size. MeBü allow for any selection procedure, so long as two assumptions hold: i) all the spurious covariates have the same random distribution of being selected, called “exchangeability”; and ii) the true-covariates are selected with higher probability. While CJSboost can satisfy the second assumption, the multi-parameter likelihood may violate the exchangeability assumption; for example, when a covariate significantly influences capture-probability but not survival, such structural correlations may make certain covariates more selectable than others. Later on, ShSa weakens these requirements through a special variant of stability selection called complementary-pairs SS.

#### Univariate vs. Multiple-Parameter Regularization

The theoretical properties derived by Bach, MeBü, and ShSa were all based on univariate least-squares regularization. Stability selection has since been used for univariate GLMs and GAMs (see Hofner et al., 2015, and citations therein). At the time of writing this article, no stability selection work has been published in a multiple parameter context, for example, using a boosted-GAMLSS model. It is unknown whether any of the theoretical properties of univariate stability selection hold for multiple-parameter regularization, or for a HMM like CJSboost. Two obvious issues arise. First, what is the effect of having different generative models for each parameter in the likelihood, and does this violate the exchangeability assumption? For example, does a *k*^th^ covariate with a significant effect in one parameter *θ*_1_ result in a biased-high estimate of 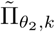 for another parameter *θ*_2_? My simulations suggest that this is not an issue and such covariates have the same null-distribution of 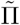 values as covariates which are spurious for both *θ*_1_ and *θ*_2_. Secondly, stability selection demands that we compute 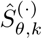 for all *reasonable* values of the regularization parameters. This is simple in univariate boosting with only one regularization parameter, but it becomes computationally unfeasible when the regularization parameter space is bivariate or trivariate (*m* and ***ν***). I have proposed a short-cut to set 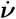 to their prediction optimized values, and then calculate 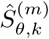 over *m* conditional on 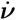. In simulations, this seems to lead to reasonable 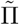 values.

#### Subsampling and Resampling

Bach used the bootstrap, whereas MeBü used subsampling at a rate of 50%, and ShSa used complementary-pairs sampling by repeatedly dividing the data into equal-halves, but acknowledged the similarity to bootstrapping. For MeBü and ShSa, the exact rate is important for deriving an upper bound on the expected number of False Discoveries (FD) in least-squares regularization. Their bounds do not apply naively to multi-parameter regularization, and so there is no reason in CJSboost to maintain their 50% subsampling rate, which otherwise has some disadvantages. For example, Schmid et al. (2012) had to subsample at a rate of 80%, and, in lieu of ShSa’s theorectical control on the FDs, they focused instead on rejecting unimportant covariates with 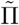 values below an arbitrary threshold *π*_thr_ *∈* (0.6, 0.9). To justify this alternative use of stability selection, Schmid et al. relied on statements by MeBü that exact values of *π*_thr_ *≫* 0.5 have little impact on the FD error rate. Bach took a different approach, and first found a theorectical region of the Lasso’s regularization parameter *λ* and sample-size, where truely influential covariates would be selected with probability *≈* 1, and spurious covariates would be selected randomly, due to the vagaries of the sampled data. Therefore, if one had multiple independent realizations of the data, then one could run the Lasso on all datasets, intersect the selection probabilities, and discard covariates *<* 0.9–1. Of course, one never has multiple independent datasets, and so Bach suggests the bootstrap to kull covariates that seem to be selected at random. In CJSboost, it is not clear whether the theorectical properties of the Bolasso hold, but I rely on research that shows how the Lasso and statistical boosting are near-equivalent estimators (Bühlmann & Yu, 2003; Efron et al., 2004). Nonetheless, the intuition behind the Bolasso bootstrap is the same: spurious covariates will have some random selection probability *«* 1. This makes SS-CJSboost crudely similar to the Bolasso, or the *adhoc* application of stability selection as in Schmid et al. (2012): we calculate inclusion probabilities and pick a high threshold to reject non-influential or insignificant covariates, in hopes of obtaining consistent model-selection.

#### Role of the Regularization Parameter

Stability selection probabilities 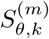 are calculated per value of a regularization parameter *m*, while inclusion probabilities 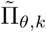 are some marginalization over *m*. MeBü used a *max* operator. ShSa suggested a *mean* operator, which results in biased 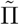 values but with much lower variance. Richardson (2010) questions whether some other integration over *m* is desirable. In simulations with CJSboost, I tried both *max* and *mean* operators, and there was considerably better separation between true and spurious covariates with the *mean* operator. Using the max operator, the overall results were very similar to Figure 7, except that the spurious covariates obtained higher 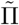 values, and there was a lot more variability among 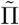 values. Also, the spurious time-as-a-categorical variables converged to *≈* 1, for both *ϕ* and *p*.

#### Inclusion InclusionProbabilities

The idea that stability selection can be used to approximate Bayesian posterior inclusion probabilities was mentioned in the Discussion and Rejoinder of MeBü by Richardson (2010) and Draper (2010). Therefore, I suggest that 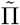 values represent interpretable end-points for a CMR analysis and can lead to correct inferences about the significance of covariates, as in Bayesian multi-model studies. Further study will be necessary to elucidate the implied prior and whether there is any meaning in the 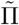 values beyond their original role as thresholding statistics. The original developers of stability selection did not espouse such a view: MeBü and ShSa use 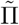 as a threshold to control FDs; Schmid et al. (2012) wished to pre-screen a high-dimensional dataset of its spurious covariates; and Bach explicitly desired a method to discard covariates and derive a consistent estimator. In other words, stability selection and 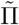 are tools to threshold one’s candidate set of covariates, and then perform estimation (but see Leeb & Pötscher, 2008). Direct interpretation of the 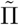 values will require further study, but the CJSboost simulations suggest that 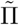 offer a fruitful means of inference about the true model.

1 In the standard AIC formula, the first term is negative. It is omitted here because I define *L* as the *negative* log-Likelihood.

2 Despite the existence of an implicit “true model”, the performance of the estimators were *not* judged on their ability to find it. Rather, the AIC and boosting are supposed to find/produce a model that minimizes the Expected negative log-Likelihood.

3 This is not to be confused with classical Null Hypothesis Tests of the marginal effect of regression coefficients.

4 After hard-thresholding, the final model may not have a unique MLE, such as as the *ϕ*(*t*)*p*(*t*) model. In such cases, one must impose constraints (such as *ϕ*_*T*–1_ = *ϕ*_*T*_) before attempting to debias the results and run the algorithm until *m* → ∞. Regularized CJSboosting does not have this problem because of shrinkage.

